# Quantitative characterization of CTLA4 trafficking and turnover using a combined *in vitro* and *in silico* approach

**DOI:** 10.1101/106898

**Authors:** Sahamoddin Khailaie, Behzad Rowshanravan, Philippe A. Robert, Lucy S. K. Walker, David M. Sansom, Michael Meyer-Hermann

## Abstract

CTLA4 is an essential negative regulator of T cell immune responses and is a key checkpoint regulating autoimmunity and anti-tumour immunity. Genetic mutations resulting in a quantitative defect in CTLA4 are associated with the development of an immune dysregulation syndrome. Endocytosis of CTLA4 is rapid and continuous with subsequent degradation or recycling. CTLA4 has two natural ligands, the surface transmembrane proteins CD80 and CD86 that are shared with the T cell co-stimulatory receptor CD28. Upon ligation with CD80/CD86, CTLA4 can remove these ligands from the opposing cells by transendocytosis. The efficiency of ligand removal is thought to be highly dependent on the processes involved in CTLA4 trafficking. With a combined *in vitro-in silico* study, we quantify the rates of CTLA4 internalization, recycling and degradation. We incorporate experimental data from cell lines and primary human T cells. Our model provides a framework for exploring the impact of altered affinity of natural ligands or therapeutic anti-CTLA4 antibodies and for predicting the effect of clinically relevant CTLA4 pathway mutations. The presented methodology for extracting trafficking rates can be transferred to the study of other transmembrane proteins.

## Introduction

In a healthy organism, the immune system has to maintain a balance between immune activation and inhibition. The decision between these two outcomes is influenced by receptors which are not specific for any antigenic stimulus. CD28 and CTLA4 are two such transmembrane receptors expressed by lymphocytes with opposing regulatory functions. These receptors bind to the same co-stimulatory ligands, CD80 and CD86, which are expressed by antigen presenting cells [1, 2]. While ligation of CD28 with co-stimulatory ligands is essential for full T cell activation and effector functions [3–5], CTLA4 inhibits excess or aberrant co-stimulation of T cells by competing for CD28 ligands and thereby preventing uncontrolled T cell activation and clonal expansion of cells specific for healthy tissues [6–8].

CTLA4 molecules are mostly observed in cytoplasmic vesicles [9–12] by the virtue of their interaction with the μ2 subunit of the clathrin adaptor protein complex AP2 [13–16]. In contrast, CD28 is present on the plasma membrane with a slow turnover rate [17]. The unusual localization of CTLA4 raises questions about the mode of CTLA4 action. For example, it is not clear how the particular distribution of CTLA4 molecules results from dynamic trafficking between cytosol and plasma membrane. The impact of various parameters affecting internalization, degradation and recycling rates all have the potential to influence the cellular distribution and function of CTLA4. Accordingly, mathematical models are useful in understanding and predicting the impact of changes in these parameters on CTLA4 localization and capacity to elicit suppressive function.

Early data suggested that CTLA4 directly inhibits the cells that express it, referred to as the cell-intrinsic pathway of immune inhibition [18,19]. Such cell-intrinsic mechanisms predict that autoimmunity arises in a CTLA4-deficient model due to uncontrolled T cell activation. Indeed, this prediction is supported by experiments. Cell-intrinsic inhibition would also imply that a chimeric model with a mixture of CTLA4-deficient and CTLA4-sufficient T cells still suffers from overactivation of CTLA4-deficient cells. However, a normal immune system with no T cell hyperactivation was found when this idea was tested experimentally [20–23]. This observation suggests CTLA4 acting through a cell-extrinsic mechanism of inhibition. Recently, it has been shown that CTLA4 molecules on T cells are able to internalize co-stimulatory molecules from the surface of APCs, a process known as transendocytosis [24]. Transendocytosis therefore represents a potential mechanism for cell-extrinsic inhibition [25].

Transendocytosis is an unusual mechanism for regulating membrane-bound ligand expression in the immune system. It allows CTLA4-expressing T cells to regulate the expression of CD80/CD86 molecules by APCs, therefore limiting CD28 signaling and activation of other T cells interacting with the ligand-depleted APCs [26]. In such a cell-extrinsic mechanism, the dynamics of CTLA4 trafficking becomes critical. Appropriate trafficking of CTLA4 and CTLA4 complexed with CD80 or CD86 molecules is required to ensure functionality. Accordingly, the efficiency of CTLA4 function is likely to be dynamically regulated by the amounts of CTLA4 delivered, the rate of removal and the time CTLA4 expressing cells spend in contact with their APC targets. In contrast, in a static model of CTLA4 expression, an efficient removal of co-stimulatory molecules from opposing cells is unlikely to be achieved. In this study, we extract CTLA4 trafficking rates from a series of experimental protocols and corresponding mathematical models in order to understand and predict the efficiency of interventions targeting CTLA4.

## Results

### General strategy

The rates of CTLA4 internalization, recycling and degradation were estimated from a series of independent experiments in which particular processes of CTLA4 trafficking were blocked or in which particular staining procedures of membrane-bound CTLA4 were applied. These experiments were replicated by corresponding mathematical models. The parameters of the model are determined by simultaneous fitting to all experimental results with a combined cost function (see *Materials and Methods - Parameter estimation*). The experimental data that we used in this study were sufficient to determine all of the CTLA4 trafficking parameters. While the parameters were identified based on all experiments, for the purpose of clarity, the different experiments are presented one by one together with their respective mathematical model.

### Ligand-independent model of CTLA4 trafficking

After synthesis, CTLA4 is transported from the Golgi to the cell surface. This transport is dependent on the enzymes phospholipase D and ADP ribosylation factor 1 [27]. Delivery of CTLA4 molecules is thought to occur via the fusion of CTLA4-containing vesicles with the plasma membrane. Interaction of the tyrosine-based motif (YVKM) in the cytoplasmic domain of CTLA4 with clathrin adaptor complex AP2 is involved in the internalization of CTLA4 from the plasma membrane to the cytoplasm [13–16, 28]. Internalized CTLA4 molecules can be recycled to the plasma membrane or degraded in lysosomal compartments [17, 29, 30], although internalized CTLA4 has also been observed in non-lysosomal, perinuclear intracellular vesicles [27, 30]. CTLA4 constitutively undergoes endocytosis independent of ligand binding, and undergoes recycling and degradation as part of its natural trafficking process [26, 28]. A detailed understanding of this trafficking itinerary is critical in understanding rates of ligand capture and ultimately the functional efficacy of CTLA4.

Based on these biological observations, a trafficking model is constructed which comprises processes of protein synthesis, recycling, internalization and degradation. This model, depicted schematically in **Figure 1A**, relies on the following assumptions: CTLA4 molecules are synthesized at a constant rate σ_*i*_. Newly synthesized CTLA4 molecules are incorporated to the plasma membrane (*R_p_(t)*) and subsequently internalized into the cytoplasm (*R_c_(t)*) with rate *k_i_*. This assumes that newly synthesized CTLA4 molecules do not directly undergo degradation in the cytoplasm, but only after exocytosis and subsequent endocytosis. Cytoplasmic CTLA4 molecules are degraded in lysosomal and non-lysosomal pathways with constant rates δ_*l*_ and δ_*n*_, respectively. Therefore, the total cytoplasmic degradation rate is δ_*c*_ = δ_*l*_ + δ_*n*_. Cytoplasmic CTLA4 molecules are exocytosed (recycled) to the plasma membrane with constant rate *k*_*r*_.

**Figure 1:**
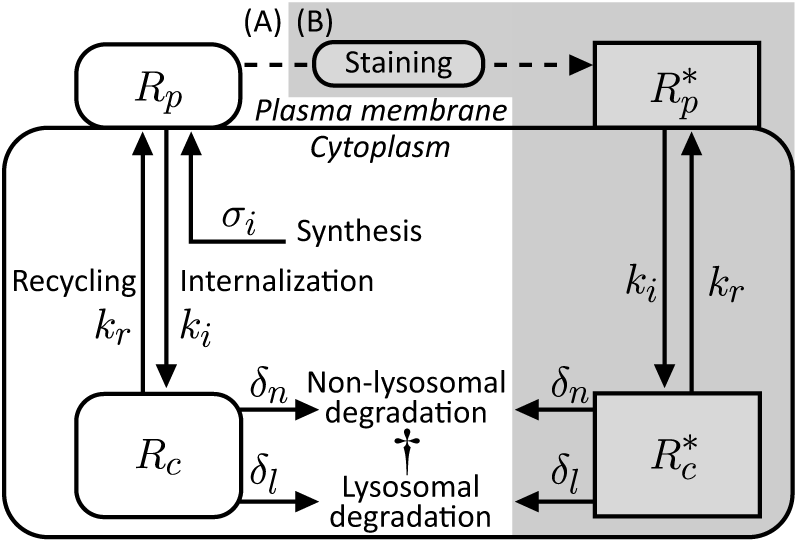
Ligand-independent model of CTLA4 trafficking. (A) Natural trafficking of CTLA4 molecules. (B) By extracellular CTLA4 staining with antibody, the CTLA4 pool is segregated into labeled (shown with grey background) and unlabeled pools. This is a simplified model of a more complex staining model described in *Materials and Methods* and shown in figure 13.

This model can be represented by the following nonhomogeneous linear system of ordinary differential equations (ODE)

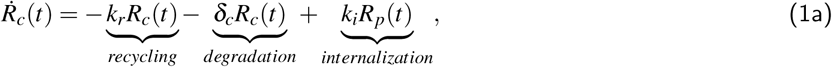

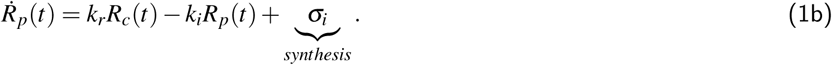

Over-dots denote time derivatives. Steady state trafficking of CTLA4 is characterised by

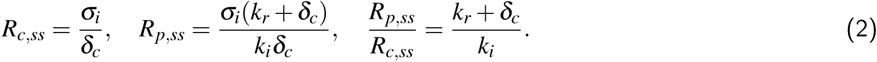

To recapitulate the experimental observation that CTLA4 molecules are mostly located in the cytoplasm, the model has to satisfy the condition *k_i_* > *k_r_* + δ*_c_* (see equation (2)). Despite the dependence of steady state CTLA4 numbers on the rate of protein synthesis, the ratio of surface to cytoplasmic CTLA4 is synthesis-independent. Further, the steady state absolute number of the cytoplasmic CTLA4 molecules is independent of the rates of internalization and recycling.

### CTLA4 staining

CTLA4 staining is based on the approach of staining surface CTLA4 with specific fluorescent antibodies. By this method, all surface molecules present on the plasma membrane are labeled at a particular time point. Subsequently, trafficking of CTLA4 is controlled by changes in the temperature of the culture. This basic principle was used with various protocols to monitor internalization and recycling of CTLA4 molecules (see subsequent result sections).

The general mathematical model for the dynamics of CTLA4 staining is described in *Materials and methods* (see **Figure 13** and equation (8)). Several modifications of this CTLA4 trafficking model (1) are needed to reflect the various staining protocols. In order to avoid unnecessary complications of the analysis by fast or too detailed processes, the model is simplified based on the following assumptions:(A) the concentration of staining antibodies in the medium is sufficiently large and does not drop by target binding (thermic bath approximation), (B)staining antibodies bind to CTLA4 molecules with a higher rate than the rate of trafficking processes (quasisteady-state approximation), (C) staining antibodies remain bound to CTLA4 molecules during the time of sample observation (negligible unbinding rate), and (D) staining antibodies do not alter the natural trafficking of CTLA4 molecules and associated rates.

The simplified model is shown in **Figure 1** and reads (see *Materials and methods - Reduction of the CTLA4 staining model*):

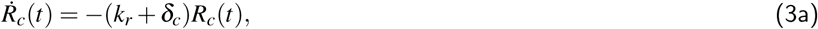

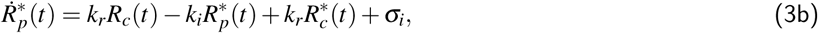

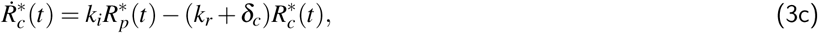

Where 
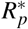
 and 
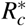
 are the numbers of labeled CTLA4 molecules on the cell surface and in the cytoplasm, respectively. Note that there is no equation for unlabeled surface CTLA4 *R_p_*(*t*), because all free surface CTLA4 molecules are labeled instantly when exposed to staining antibodies in the medium (assumption B). Before starting the staining experiment, the number of CTLA4 molecules is assumed at equilibrium given in equation (2). During the staining process at 37°C, the unlabeled pool of cytosolic CTLA4 molecules becomes exhausted with a half-life of *t*_1/2_ = ln(2)=(*k*_*r*_ + δ_*c*_) ≈ 27 minutes (see equation (17a)). Therefore, the rates of recycling and degradation dictate the duration of the staining process required to label the majority of the cellular CTLA4 molecules.

In experiments with time-limited staining followed by removal of excess antibodies, recycled and newly synthesized molecules cannot be labeled further. Then, the dynamics of the labeled CTLA4 molecules in equation (3b) is replaced by

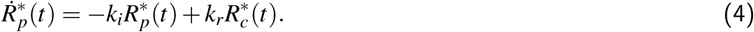

The kinetics of different pools of CTLA4 molecules during persistent staining is shown in **Figure 2**. Due to instant labeling of surface and newly synthesized CTLA4 molecules as well as continued exocytosis, the amount of labeled CTLA4 molecules on the cell surface starts and remains at its equilibrium value. Unlabeled cytoplasmic CTLA4 molecules become exhausted over time since no free CTLA4 molecules can be found on the plasma membrane. The number of the labeled CTLA4 molecules saturates when the cytoplasmic unlabeled CTLA4 molecules are exhausted.

**Figure 2:**
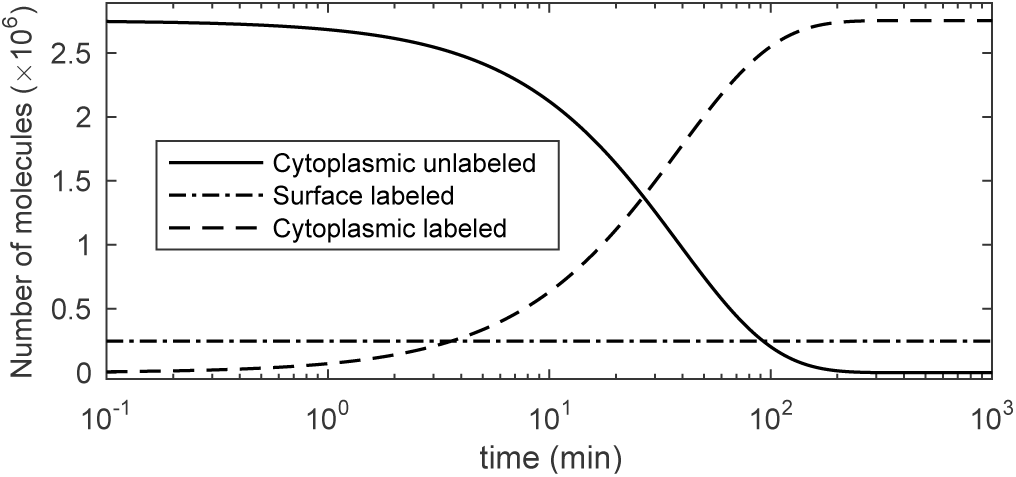
CTLA4 staining simulation. Equation (3) was solved analytically and the solution, given in equation (17), was plotted.

### CTLA4 internalization

To observe the internalization of surface CTLA4 molecules, we used a CTLA4 internalization assay described in *Materials and methods*. In this experiment, CTLA4-expressing CHO cells were labeled at 4°C with a fluorescently conjugated anti-CTLA4 antibody (PE-CTLA4). The excess antibody is washed off and the temperature is raised to 37°C for various durations to allow for CTLA4 trafficking. After this period, a secondary antibody (Alexa fluor 647) is applied at 4°C. In this setting, the initial amount of surface CTLA4 is quantified by PE and the amount still at the surface after a period of trafficking is quantified by Alexa 647. The data of this experiment are shown in **Figure 3A**.

**Figure 3:**
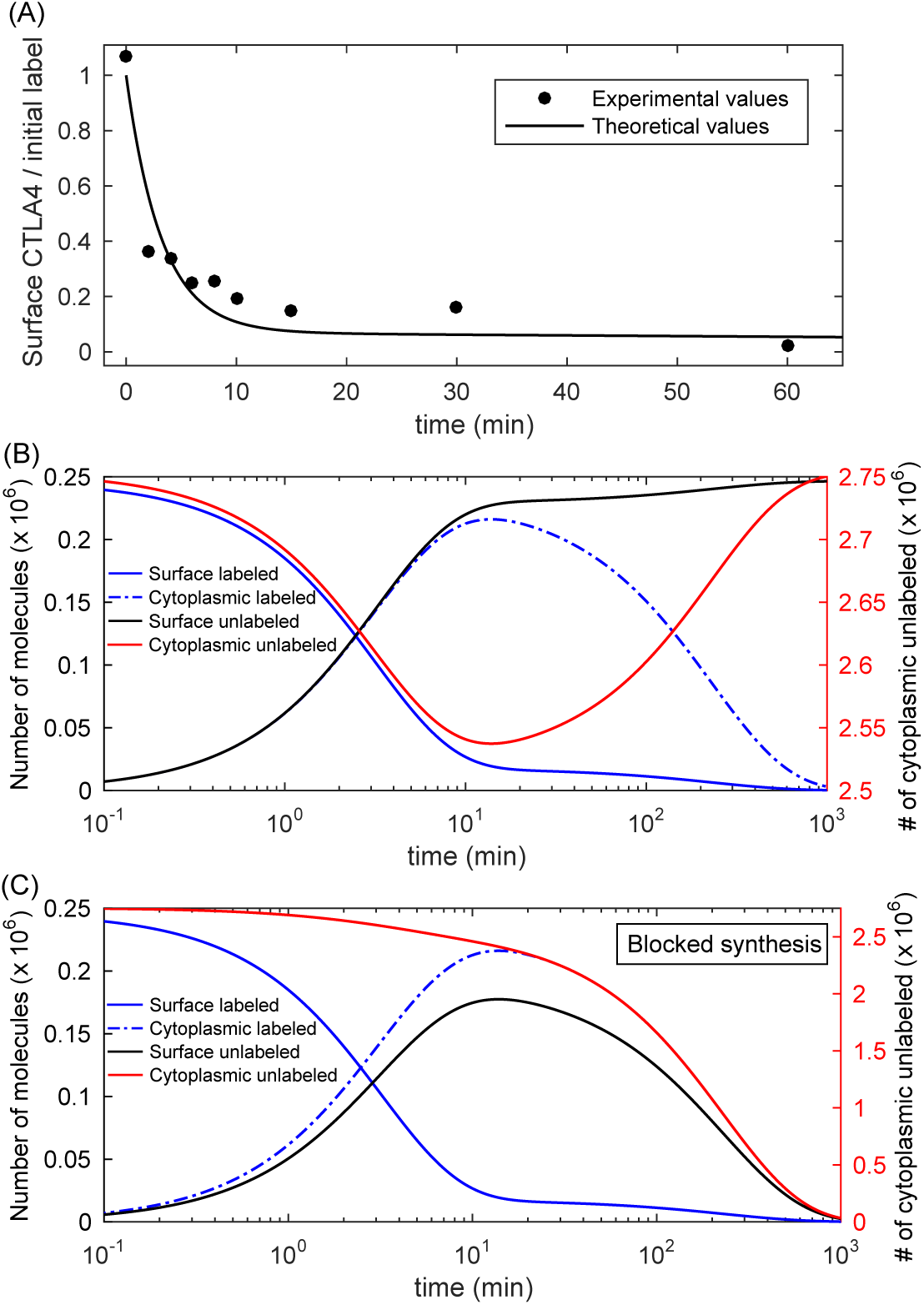
*In silico* and *in vitro* internalization data. (A) The fraction of initially labeled CTLA4 molecules detected on the cell surface over time is shown. Experimental data points are scaled with the factor 1 = *S*_int_ (see Table 1). (B) The kinetics of all different CTLA4 pools in the model. Unlabeled CTLA4 follows equation (1). Unlabeled cytoplasmic CTLA4 molecules (*R_c_(t)*) drop transiently (right y-axis). (C) An *in silico* internalization experiment with a block of CTLA4 synthesis. Both unlabeled CTLA4 pools clear over time; however, the surface unlabeled pool transiently reaches 72% of its steady state value in normal cells.

**Table 1.** Ligand-independent CTLA4 trafficking parameters

| Parameter | Description | Dimension | Estimated value |
| --- | --- | --- | --- |
| $\sigma_i$ | Rate of CTLA4 synthesis | $\# \text{ min}^{-1}$ | 12830 |
| $k_i$ | Rate of CTLA4 internalization | $\text{min}^{-1}$ | 0.291 |
| $k_r$ | Rate of CTLA4 recycling | $\text{min}^{-1}$ | 0.021 |
| $\delta_n$ | Rate of non-lysosomal degradation | $\text{min}^{-1}$ | 0.0002 |
| $\delta_l$ | Rate of lysosomal degradation | $\text{min}^{-1}$ | 0.0044 |
| $\delta_c$ | Rate of cytoplasmic degradation | $\text{min}^{-1}$ | $\delta_l + \delta_n$ |
| $S_{\text{int}}$ | Scaling factor in internalization model; eq. (20) | - | 0.933 |
| $S_{\text{rc}}$ | Scaling factor in recycling model; eq. (37) | - | 0.996 |
| $S_{\text{NH}_4\text{Cl}}$ | Scaling factor in degradation model (+NH<sub>4</sub>Cl); eq. (28a) | - | 0.964 |
| $S_{\text{CTR}}$ | Scaling factor in degradation model (-NH<sub>4</sub>Cl); eq. (28b) | - | 1.009 |
| $\Delta t_{\text{rc}}$ | Shifting time in recycling data; eq. (37) | min | 5.75 |
| $R$ | Radius of CHO cell | $\mu\text{m}$ | 7; fixed |
| $z_s$ | Stack size of confocal microscope | $\mu\text{m}$ | 1; fixed |

In the model, the amount of surface CTLA4 molecules before starting the staining process is known from the steady state solution of the CTLA4 trafficking model, given in equation (2). This amount is assumed to be fully labeled on ice at *t* = 0, leaving no unlabeled molecules on the cell surface. The solution of the dynamic equations provides the time course of the amount of different CTLA4 pools (see *Materials and methods*, equation (18)). Surface labeled CTLA4 molecules vanish over time monotonically, becoming cytoplasmic due to internalization (**Figure 3B**), while the cytoplasmic labeled CTLA4 pool vanishes after a transient increase due to degradation. Newly synthesized and recycled unlabeled CTLA4 molecules replace the labeled pool over time.

An *in silico* internalization experiment with an additional blocking of CTLA4 synthesis (σ_*i*_ = 0) shows that unlabeled CTLA4 molecules on the plasma membrane are cleared only after a transient recovery to 72% of its pre-experiment (normal steady state) value (see **Figure 3C**). This recovery results from the recycling of the cytoplasmic CTLA4, and suggests that, when CTLA4 synthesis is suppressed, the large amount of CTLA4 stored in the cytoplasm can temporarily compensate the loss of surface CTLA4 expression.

### Block of CTLA4 synthesis

To quantify the contribution of protein synthesis to CTLA4 homeostasis, we utilized the data from an experiment in which CTLA4-expressing CHO cells were treated with Cycloheximide (CHX) to block protein synthesis. In this experiment, CHO cells were treated for 3 hours with CHX, which resulted in an approximately 55% reduction of the total number of CTLA4 molecules [28].

In the model, we assume that CHX treatment of cells completely blocks protein synthesis. This is achieved by removing the process of synthesis in the CTLA4 trafficking model in equation (1), i.e. imposing *σ_i_* = 0. The time evolution of surface and cytoplasmic CTLA4 in the presence of CHX is calculated (see *Materials and methods*, equation (21)) and depicted in **Figure 4A**. At early time points, surface CTLA4 molecules quickly drop due to fast internalization and lack of compensation by newly synthesized molecules. Recycling partially compensates surface CTLA4 loss until CTLA4 molecules vanish by degradation in the cytoplasm. The half life of surface, cytoplasmic and total CTLA4 molecules are 121, 163 and 160 minutes, respectively. Surface CTLA4 drops by 10.5% (≈26000 molecules) within 3 minutes while cytoplasmic CTLA4 drops by only 0.5% (≈14000 molecules) during this time. As expected, CTLA4 synthesis is important for maintaining the homeostatic number of CTLA4 molecules.

**Figure 4:**
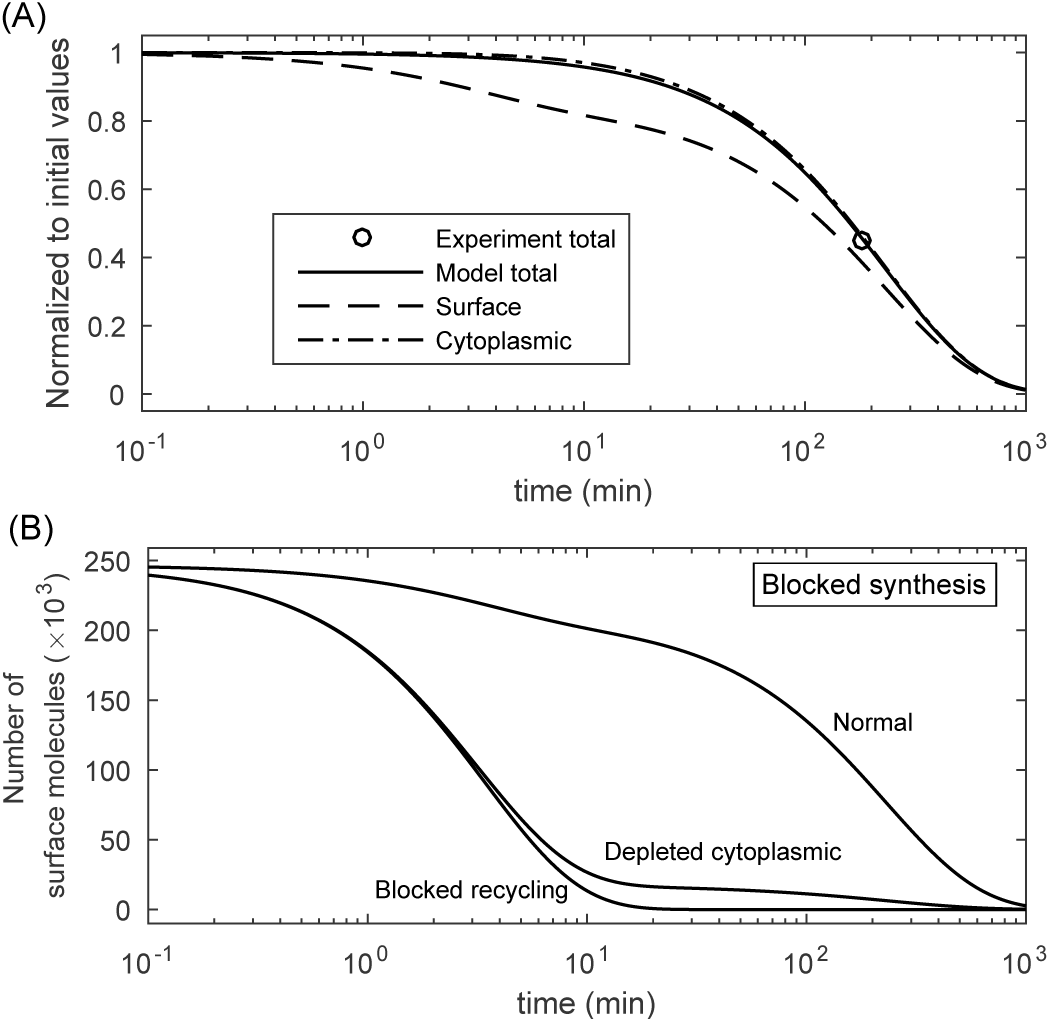
Block of CTLA4 synthesis. (A) *In silico* versus *in vitro*. A single experimental data point was used in the parameter estimation process in order to fit the theoretical value in equation (24) to the measurement. Each curve is normalized to its own initial value. CTLA4 molecules in the cytoplasm can transiently compensate CTLA4 loss on the plasma membrane after the block of synthesis. However, both pools become exhausted with extended exposure to CHX. (B) *In silico* experiment of blocked synthesis with additional depletion of cytoplasmic CTLA4 (*R_c,0_* = 0) or block of recycling (*k_r_* = 0). These curves are obtained by numerical solution of the trafficking model in equation (1). In the absence of cytoplasmic CTLA4 or of the recycling process, the surface CTLA4 clears quickly. When the recycling process is blocked, clearing of the surface CTLA4 is affected only by internalization process with the half-life of 2.38 minutes.

Next, we asked how important internalization would be for CTLA4 surface expression. In an *in silico* experiment, in which the block of CTLA4 synthesis is complemented with a complete depletion of the cytoplasmic CTLA4 (by forcing a zero initial value) or with a complete block of recycling (*k_r_* = 0), the surface CTLA4 molecules clears within a few minutes in comparison to 2 hours under normal conditions (see **Figure 4B**). This suggests that recycling is essential for the stability of surface CTLA4 expression.

### Block of lysosomal degradation

To quantify the impact of lysosomal degradation on CTLA4 trafficking, we took advantage of data from an experiment in which CTLA4 kinetics were measured in presence or absence of ammonium chloride (NH_4_Cl). NH_4_Cl inhibits lysosomal degradation through lysosomal pH neutralization. In this experiment, cells were initially pulsed with a labeling antibody (anti-CTLA4 PE) at 37°C for 60 minutes. After washing, degradation of CTLA4 molecules was quantified by flow cytometry (see **Figure 5A**) [28].

**Figure 5:**
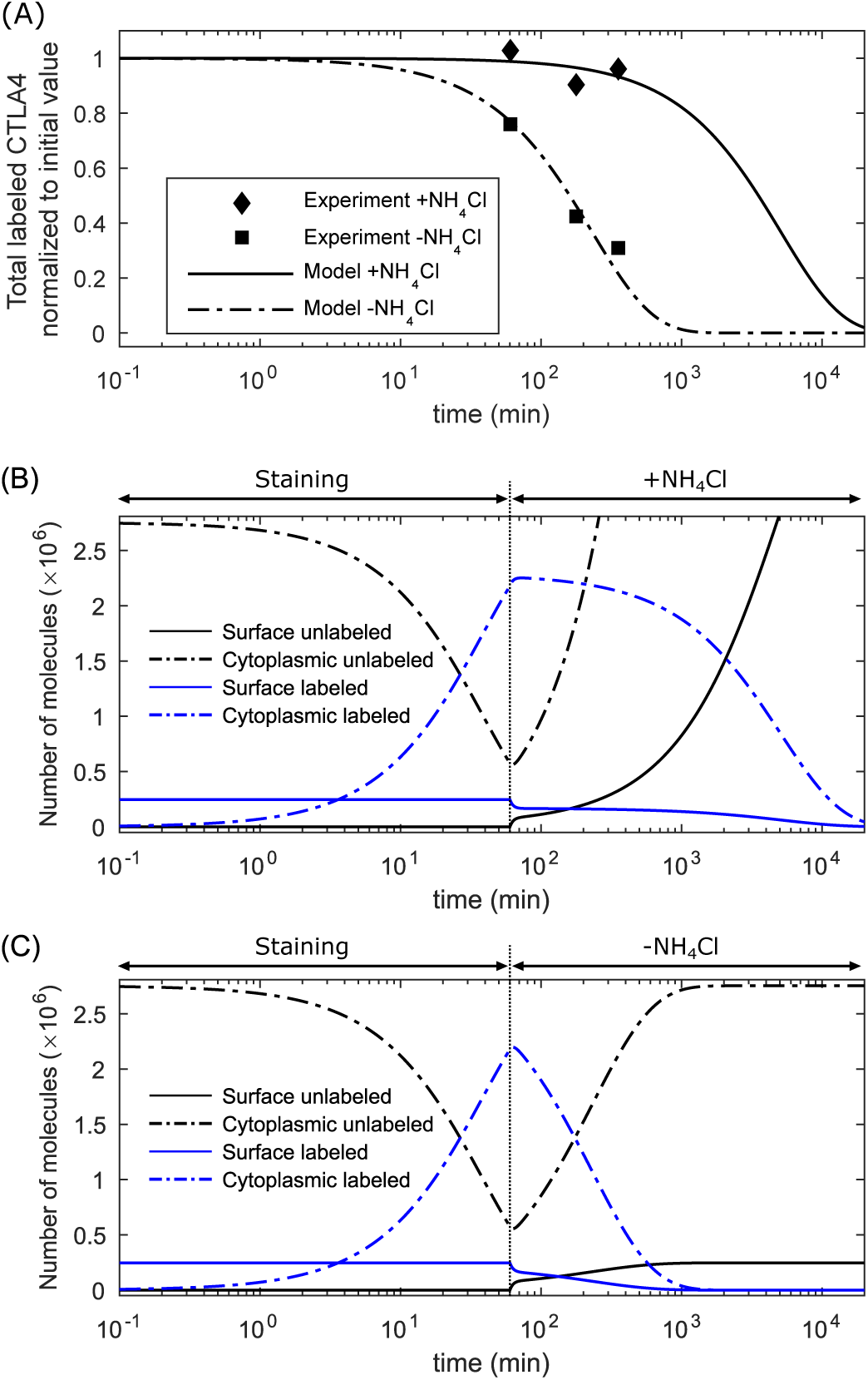
Block of lysosomal-degradation. (A) *In silico* versus *in vitro* results in two different conditions. Experimental data points are scaled with factors 1/*S*_NH_4_Cl_ and 1=*S*_CTR_ for blocked lysosomal degradation and control experiments, respectively (see Table 1). (B) *In silico* prediction of different CTLA4 pools for blocked lysosomal degradation and (C) normal condition.

To reconstruct this experiment *in silico*, we used the staining model in equation (3) and calculated the amount of labeled CTLA4 molecules after 60 minutes. Starting from this value as initial condition, the time evolution of the CTLA4 pools was obtained (see *Materials and Methods*, equation (25)). *In silico* simulation of this experiment for both stages, initial staining and follow up kinetics, are shown in **Figure 5B-C** in the presence or absence of NH_4_Cl, respectively. It was assumed that NH_4_Cl does not impact on the nonlysosomal degradation pathway (with rate δ_*n*_). A comparison of both figures shows the differences in the kinetics of labeled CTLA4 molecules: The rate of degradation decreased by ≈96% in the presence of the lysosomal inhibitor. The half-life of total labeled CTLA4 molecules increases from approximately 2.7 hours to 59 hours by adding the lysosomal inhibitor. After the initial decrease of the number of unlabeled CTLA4 molecules during the staining step, it increases to its steady state value or higher in the absence or presence of the lysosomal inhibitor, respectively. This increase is due to the constant synthesis of CTLA4.

In an *in silico* experiment, in which the lysosomal degradation is blocked (δ_*l*_ = 0) for 3 hours, the number of unlabeled CTLA4 molecules increases ≈1.72-fold, which is close to the experimentally observed 2-fold increase of CTLA4 level in the presence of NH_4_Cl [28]. According to equation (2) (see *R_p,ss_*/*R_c,ss_*), by reducing the cytoplasmic degradation (δ_*c*_), the fraction of the total CTLA4 molecules that appears on the cell surface reduces.

These results suggest that the lysosomal degradation of CTLA4 is a fast process, regulates the homeostatic number of CTLA4 molecules, and, when blocked, shifts the steady state subcellular distribution of CTLA4 towards the cytoplasm.

### CTLA4 recycling

To quantify CTLA4 recycling, we used a two-step staining experiment. Two steps were required to label a subset of CTLA4 molecules that were, at least once, on the plasma membrane, and then to detect a subset of these molecules that recycled from the cytoplasm back to the plasma membrane. In the first step, CTLA4-expressing CHO cells were incubated with Alexa488-conjugated anti-CTLA4 (a green antibody) for 60 minutes at 37°C. Then, cells were washed and surface CTLA4 molecules were blocked at 4°C with unconjugated anti-human IgG, a secondary antibody without any color. This excluded those molecules that remained on the surface and were green-labeled from binding to a secondary antibody. After washing, cells were incubated with Alexa555-conjugated anti-human IgG (a red secondary antibody) at 37°C. The red antibody will only bind those green-labeled CTLA4 molecules that recycled back to the plasma membrane. The red-fluorescent intensity was quantified with confocal microscopy reflecting the time evolution of the labeled subset of recycled CTLA4. Experimental data are shown in **Figure 6A** [28].

The corresponding *in silico* experiment encapsulates the two staining steps. At first, the amount of green-labeled CTLA4 molecules stained on the cell surface and subsequently internalized within 60 minutes is calculated from the staining model in equation (3). During the CTLA4 block on ice, we assumed that all green-labeled molecules on the cell surface are blocked (denoted by green-blocked in **Figure 6B**), and undergo natural trafficking at normal temperature. In the model, application of the red secondary antibody labels only those CTLA4 molecules that recycle back from the cytosol and do not belong to the blocked subset. This defines a green-red labeled pool of molecules.

The recycling model is described in *Materials and Methods - CTLA4 recycling model*. Model predictions for the kinetics of different CTLA4 pools during the entire experiment (see equation (30) for the labeled quantities in the second staining step) are shown in **Figure 6B**.

**Figure 6:**
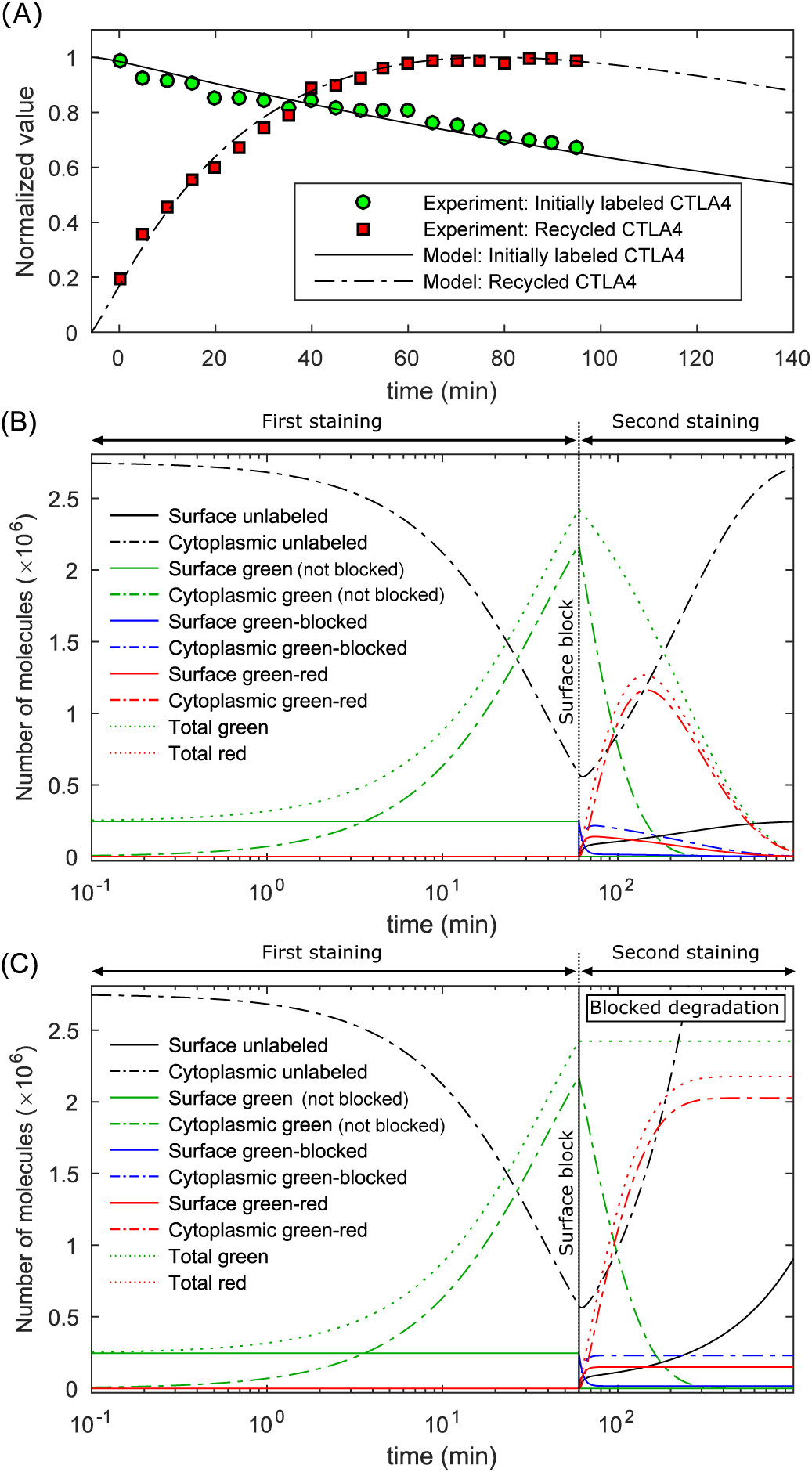
Simulation of the recycling experiment. (A) *In vitro* and *in silico* amount of green- and red-labeled CTLA4 molecules. (B) Model prediction of different CTLA4 pools during the staining procedures. The vertical line at t=60 minutes indicates the block of green-labeled CTLA4 and the starting time of the second staining step. (C) *In silico* recycling experiment with additional block of CTLA4 degradation (δ_c_ = 0) in the second staining step. The clearance of cytoplasmic green-labeled (not blocked) CTLA4 (dash-dotted green curve) in the second staining step is only affected by the recycling process with a half-life of ≈32 minutes.

According to the modeling results, in the second staining step, green-blocked molecules clear monotonically from the cell surface by internalization, while the cytoplasmic green-blocked molecules vanish after a transient increase, reaching 88% of total initially blocked CTLA4 molecules after 14 minutes (t=74 minutes) of trafficking. Such a transient increase is also observable for green-red labeled molecules, which originate from the green-labeled cytoplasmic pool. The amount of green-red labeled molecules on the cell surface reaches its maximum value (at t=74 minutes) faster than the cytoplasmic pool (at t=146 minutes), since the labeling is only possible by passing through the cell surface first. The green‑ and red-labeled CTLA4 molecules clear over time and are replaced with the unlabeled CTLA4 molecules.

In an *in silico* recycling experiment, in which the CTLA4 degradation is blocked (δ_*c*_ = 0) in the second staining step, the clearing of the cytoplasmic green-labeled (not blocked) CTLA4 is only affected by the recycling process with a half-life of ≈32 minutes (see **Figure 6C**).

These results suggest that recycling of the cytoplasmic CTLA4 occurs very fast, but significantly slower than internalization (*t*_1/2_ = 2.38 minutes, see **Figure 4B**).

### Estimating the rate of protein synthesis

The rate of protein synthesis could not be determined because the models were based on relative counts independent of the CTLA4 synthesis rate. From the model in **Figure 1A** we know that CTLA4 synthesis is related to absolute counts of CTLA4. Therefore, we did a molecular counting experiment (see *Materials and methods - CTLA4 molecular counting*). For a typical virally transduced CHO cell line, we estimated 3 × 10^6^ cellular CTLA4-WT per CHO cell, which reflects the total steady state number of CTLA4 molecules. By using this value and the fitted rates of internalization, recycling and degradation, we obtained the rate of protein synthesis from equation (2).

### Subcellular distribution of CTLA4 molecules

The ratio of surface to cytoplasmic CTLA4 in steady state condition, given in equation (2), is independent of the rate of CTLA4 synthesis. However, by changing this rate, the trafficking model deviates from its equilibrium state, and the absolute number of CTLA4 molecules changes to a new steady state value. This transition induces a transient change in the subcellular distribution of CTLA4. By increasing or decreasing the rate of CTLA4 synthesis, the fraction of CTLA4 molecules on the cell surface transiently increases or decreases, respectively. This distribution converges back to its original value after a sufficiently long time. **Figure 7A** shows how a sharp change in the rate of protein synthesis transiently deviates the subcellular distribution. A smooth change in the rate of protein synthesis has less impact on the distribution of CTLA4 molecules (**Figure 7B**).

These results suggest that the subcellular distribution of CTLA4 can be transiently shifted towards the cell surface by upregulation of CTLA4 synthesis.

**Figure 7:**
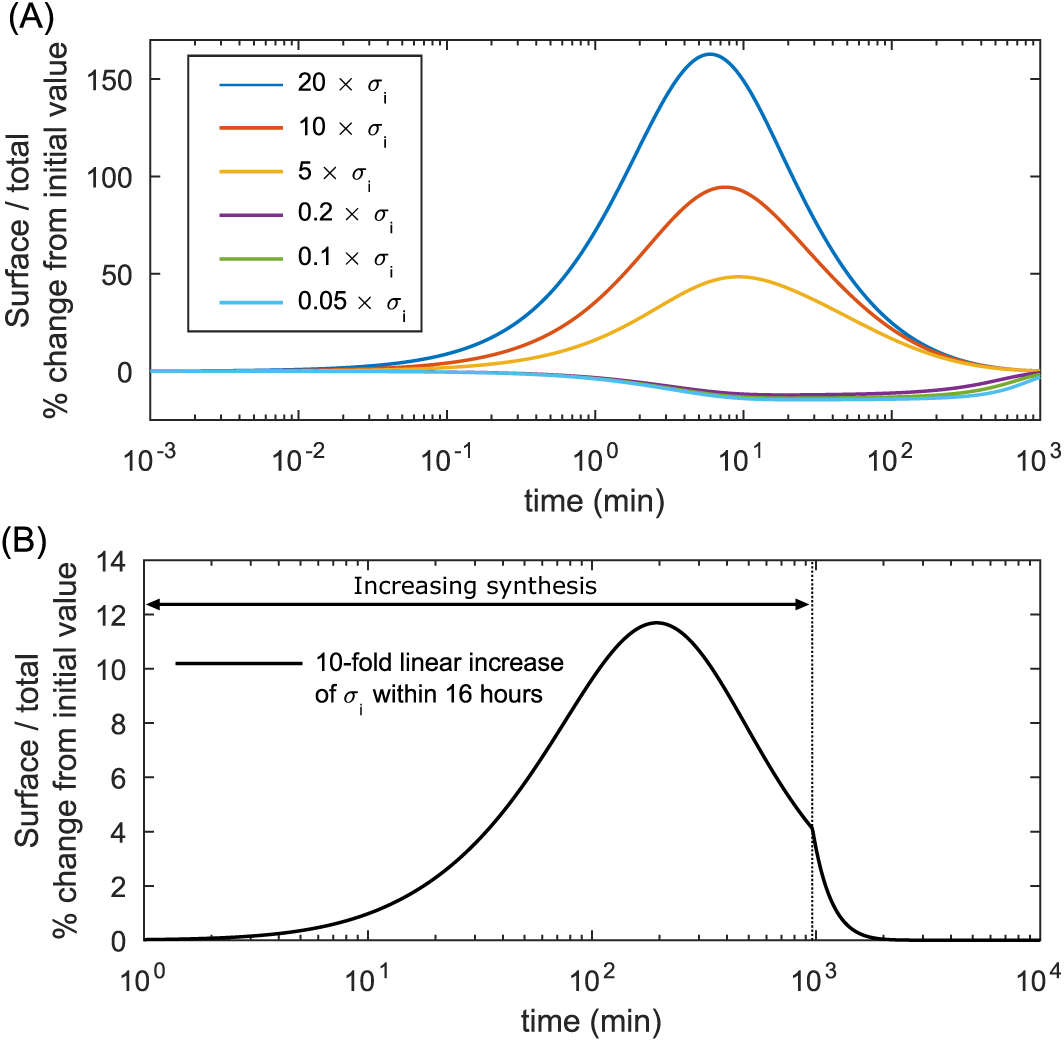
CTLA4 subcellular distribution changes transiently upon variation in CTLA4 synthesis. The rate of protein synthesis (σ_*i*_) is varied (A) sharply and (B) smoothly from its original value, and the dynamic change of the fraction of CTLA4 molecules expressed on the cell surface are obtained. After a transient change, the fraction converges to its original value after longer times.

### Amount of degraded CTLA4:antibody complex

The CTLA4 staining model in equation (3) relies on the assumption that abundant labeling antibodies remain bound to and degrade along with the bound CTLA4 molecules. Theoretically, this is a good approximation for antibodies with high affinity (high on-rate and low off-rate) to CTLA4 molecules. Similar to labeling antibodies, monoclonal antibodies that bind to CTLA4 molecules with a very high affinity do not dissociate until CTLA4:antibody complexes get degraded. Upon almost instantaneous binding to CTLA4, bound antibodies are out of the active pool in the medium.

In the context of anti-CTLA4 therapy, this could result in a significant target-mediated drug disposition, with a significant proportion of the drug (relative to dose) bound with high affinity to its pharmacological target. This interaction would be reflected in the pharmacokinetic properties of the drug. We investigated the dynamic amount of the antibody (drug) degraded along with CTLA4 molecules. The rate of antibody degradation depends on the amount of CTLA4:antibody complexes in the cytoplasm 
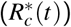
 at any given time. The total amount of degraded antibodies, denoted by *D*(*t*), can be obtained from the CTLA4 staining model in equation (3)

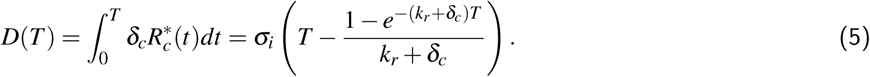

After a sufficiently long exposure to antibodies, the exponential term in *D*(*T*) converges to zero and the amount of degraded antibodies becomes linear with respect to time *T*. According to the estimated parameters of CTLA4 trafficking, it takes 4.5 hours to degrade an amount of antibodies equal to the total amount of CTLA4 molecules (see **Figure 8A**).

**Figure 8:**
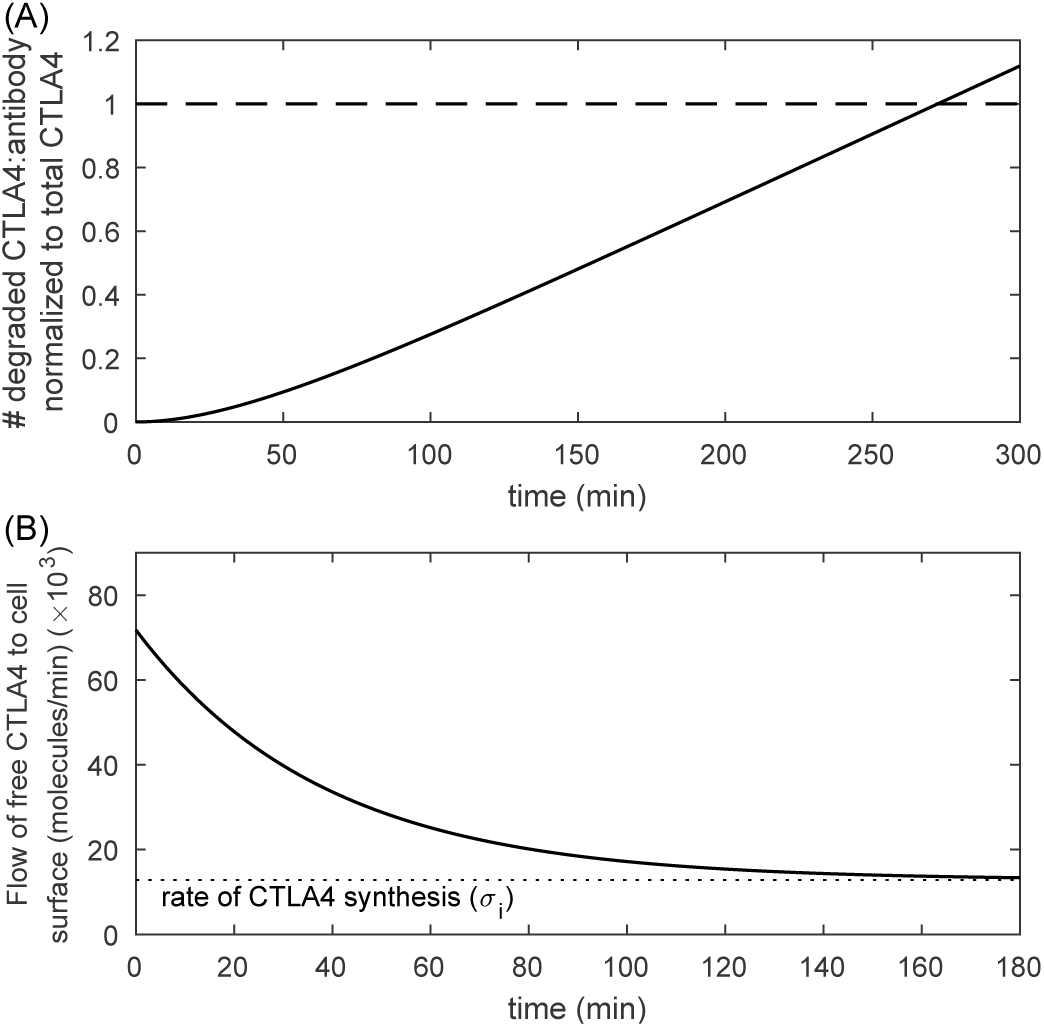
Antibody degradation and free CTLA4 flow. (A) The amount of degraded antibodies is obtained from equation (5). This amount is normalized to the steady state number of CTLA4 molecules (*R_p,ss_* + *R_c,ss_*) given in equation (2). Therefore, it is independent of the value for the rate of CTLA4 synthesis. (B) The flow of free CTLA4 molecules to the cell surface is obtained from equation (6). This flow converges to the rate of CTLA4 synthesis by clearing the pool of cytoplasmic free CTLA4 molecules. The analytic value of *R_c_(t)* is given in equation (17a).

The rate of antibody removal from the medium to the cytoplasm, in addition to the rate of internalization, also depends on the flow of free CTLA4 molecules to the cell surface (*F_cs_*), which is provided by CTLA4 synthesis and recycling of the cytoplasmic free CTLA4 molecules

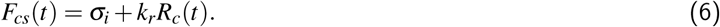

Upon exposure to antibodies, the free cytoplasmic pool of CTLA4 molecules clears over time (*R_c_*(*t*) → 0) and reduces by 98% within 2.5 hours (see **Figure 2**). Consequently, the flow of free CTLA4 molecules to the cell surface is reduced and limited to the newly synthesized CTLA4 molecules (*F_cs_*(*t*) → σ_*i*_) (see **Figure 8B**).

These results suggest that the rate of antibody passage from the medium into the cytoplasm is highest at the time of exposure to the antibody when there are still many free CTLA4 molecules in the cytoplasm, whereas the frequency of antibody degradation per time 
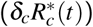
 increases when the free CTLA4 molecules in the cytoplasm get exhausted.

### The impact of binding affinity on ligand uptake

To investigate the uptake kinetics of physiological ligands of CTLA4, namely natural and drug ligands, we employed a ligand uptake model that explicitly considers CTLA4:ligand binding reactions. We relaxed the previous assumption in the staining model (equation (3)) that ligands are abundantly available in the extracellular environment (assumption A). It is assumed that modeled ligands react with surface CTLA4 with constant on-rate (*k*_on_) and off-rate (*k*_off_). The complexes can be internalized and degraded by the rate of CTLA4 internalization (*k*_*i*_) and degradation (δ_*c*_), respectively. Free modeled ligands appear in the cytoplasm via unbinding of the cytoplasmic complexes to the reactants with the rate *k*_off_. We assumed that free ligands cannot bind again to free CTLA4 in the cytoplasm, before they are degraded by a rate equal to the rate of CTLA4 degradation. Since free ligands in the cytoplasm do not interact with other trafficking molecules in the model, the absolute value of their degradation rate does not impact onto extracellular ligand uptake. For simplicity, we assumed the same rate as for the degradation of CTLA4:ligand complexes.

After unbinding, free CTLA4 molecules directly join other cytoplasmic free CTLA4 molecules (CTLA4 recovery), which subsequently can be recycled or degraded. The model is schematically depicted in **Figure 9** and reads:

**Figure 9:**
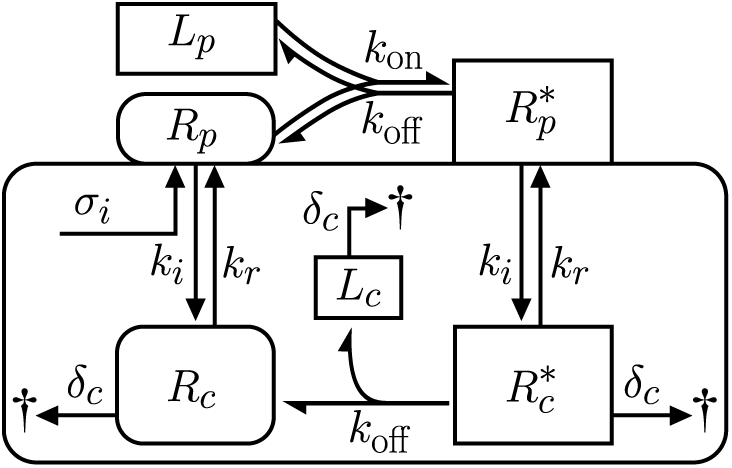
Schematic of binding reaction and trafficking of CTLA4 and modeled ligand with limited extracellular concentration.

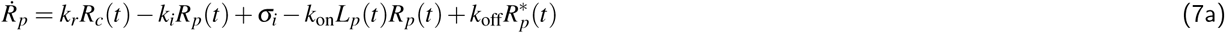

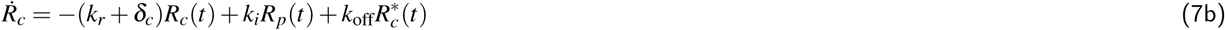

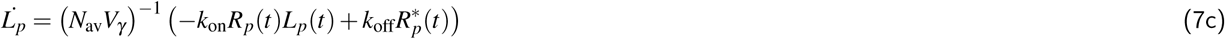

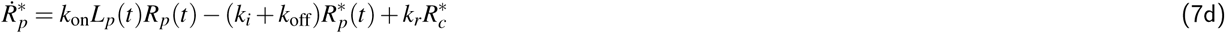

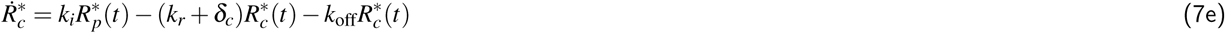

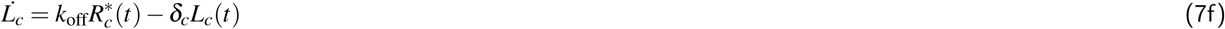

Where 
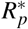
 and 
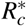
 are surface and cytoplasmic CTLA4:ligand complexes, respectively. *L*_*p*_ is the concentration of extracellular ligands, *L*_*c*_ is the number of free intracellular ligands. *N*_av_ is Avogadro’s number and *V*_γ_ is the volume of the extracellular medium per cell. We performed an *in silico* cell culture experiment with a cell count of 10^6^ cells/ml. This gives *V*_γ_ = 10 ^-9^ Liters/cell.

To investigate the role of ligand binding affinities in the kinetics of ligand uptake, equation (7) is solved numerically for ligands with different on-rates (fixed off-rates) or off-rates (fixed on-rates). The time at which half of the initial extracellular ligand concentration was depleted is monitored for different initial ligand concentrations (see **Figure 10A-B**). These results indicate how fast modeled ligands can be depleted from the extracellular environment via CTLA4 trafficking.

According to **Figure 10A**, the uptake of the extracellular ligands with higher on-rates (higher affinities) is faster. In contrast, this process is optimally fast only for an intermediate range of off-rate values (see **Figure 10B**). For low off-rates (high affinity ligands), uptake is not efficient since the recovery of CTLA4 from internalized CTLA4:ligand complex and their subsequent recycling is not efficient. For high off-rates (low affinity ligands), CTLA4:ligand complexes on the cell surface are not stable and complex internalization is limited by complex concentration, which results in an inefficient ligand uptake. We conclude that ligand removal from the medium is monotonic in dependence on on-rates but exhibits a maximum in dependence on off-rates.

To evaluate the capacity of modeled ligands to occupy free CTLA4 molecules on the plasma membrane and in the cytoplasm, the maximum reduction of free CTLA4 is obtained during the process of ligand uptake (see **Figure 10C-D**). The degree of CTLA4 occupancy on the plasma membrane depends on both, extracellular ligand concentration and ligand binding affinity (see **Figure 10C**). In the cytoplasm, it only depends on the ligand binding affinity when sufficient extracellular ligands are provided (see **Figure 10D**).

Natural ligands CD80/CD86 (dotted vertical lines in **Figure 10C-D**) strongly impact on CTLA4 occupancy on the cell surface, but have only a minor influence in the cytoplasm. This allows the cytoplasmic free CTLA4 pool to keep the flow of the free CTLA4 to the cell surface, and consequently the ligand uptake process close to its maximum value.

**Figure 10:**
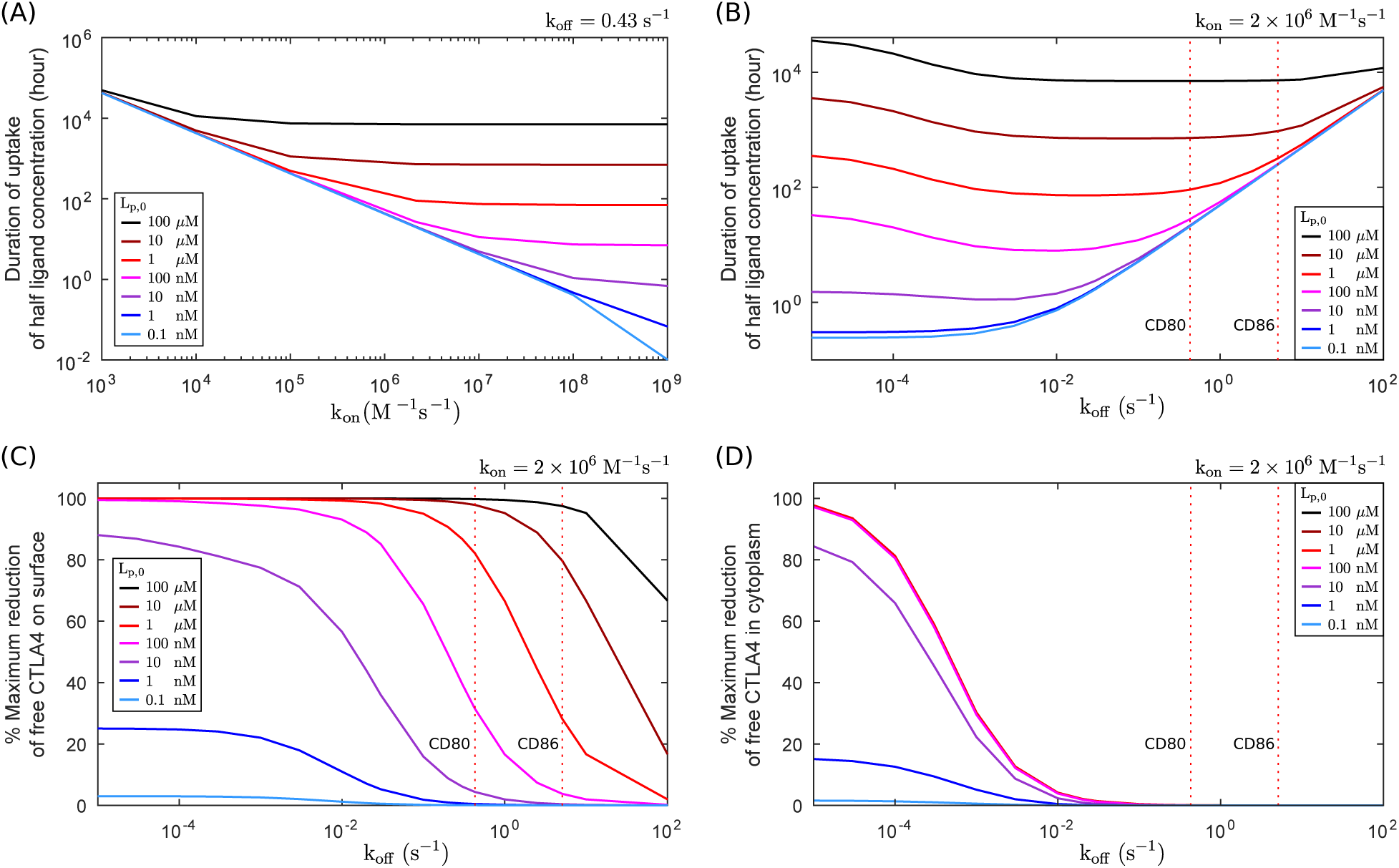
Extracellular ligand uptake and CTLA4 occupancy. The kinetics of ligand uptake is obtained numerically from model (7) for ligands with different binding affinities. Affinities are varied by changing either on-rate (k_on_), or off-rate (k_off_) of CTLA4-ligand reaction. Duration of uptaking half ligand concentration versus (A) on-rate (k_on_) or (B) off-rate (k_off_) are obtained for various initial extracellular ligand concentration. (C,D) The maximum reduction of free CTLA4 (or the degree of CTLA4 occupancy) resulted from binding to ligand is obtained for different off-rates. This reduction is shown for (C) free CTLA4 pool on the plasma membrane and (D) in the cytoplasm.

### Reduction of CTLA4 level by alterations in CTLA4 recycling and degradation

Lipopolysaccharide-responsive and beige-like anchor protein (LRBA) is an intracellular protein, which colocalizes with CTLA4 in endosomal vesicles. Patients with LRBA mutations exhibit a significant reduction in CTLA4 expression [31]. It has been suggested that the interaction with LRBA in the endosomes rescues CTLA4 from lysosomal degradation, such that LRBA deficiency results in higher degradation and lower recycling of CTLA4 [31,32]. To test this idea with the mathematical model, we checked the existence of new rates for recycling (*k*_*r*,LRBA_) and lysosomal degradation (δ_*l*,LRBA_) that represent CTLA4 trafficking in LRBA deficient cells, such that the experimentally observed reduction in CTLA4 expression can be reproduced.

Normal and LRBA-deficient human Tregs were stimulated for 16 hours with CD3/CD28 beads, with and without lysosomal inhibitor Bafilomycin A (BafA), and CTLA4 expression was measured (**Table 2**). We considered three different constraints for finding *k*_*r*,LRBA_ and δ_*l*,LRBA_, namely the ratio of total CTLA4 of LRBA-deficient cells to control cells in the presence and absence of BafA, and fold increase of CTLA4 in LRBA-deficient cells after treatment with BafA.

**Table 2.** Measurements of total CTLA4 molecules (MFI) in human Tregs.

| Condition | Control | LRBA-deficient |
| --- | --- | --- |
| Stimulation | 30000 | 15000 |
| Stimulation + BafA (16h) | 60000 | 45000 |

For given recycling rates *k*_*r*,LRBA_, the data-set in **Table 2** determined the LRBA-specific lysosomal degradataion rate δ_*l*,LRBA_ (see **Figure 11A**). An increased lysosomal degradation (by ≈1.7-fold) can explain the two-fold reduction observed in the total CTLA4 level of LRBA-deficient cells. A complete block of the recycling process alone can reduce the total CTLA4 level by only 7% (not shown). Therefore, the total CTLA4 level is not sensitive to the recycling impairment (reduction in the rate). However, the recycling process directly influences the number of surface CTLA4, and the degree of its impairment influences the kinetics for the uptake of ligands by the cells (see **Figure 11B**).

These results suggest that a decrease in recycling alone cannot account for the total CTLA4 reduction observed in LRBA-deficient cells, but it can affect the kinetics of ligand uptake and by that, impair the cell-extrinsic inhibitory function of CTLA4.

**Figure 11:**
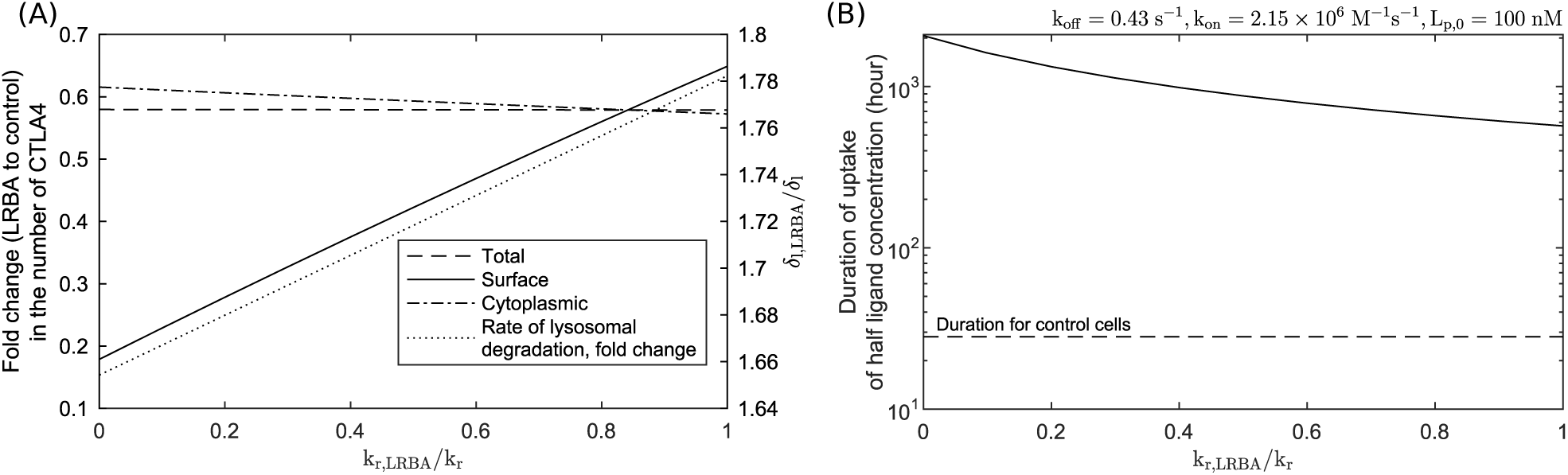
Alterations in recycling and degradation processes. (A) The rate of recycling (*k*_r,LRBA_) is fixed to different values, and the rate of lysosomal degradation (δ_l,LRBA_) is estimated such that the data constraints of LRBA-deficient and control cells in table 2 are satisfied. Fold change in the number of CTLA4 molecules (LRBA to control) is shown for total, cytoplasmic and surface CTLA4 molecules (left y-axis). Fold change in the rate of lysosomal degradation (LRBA to control) is shown (right y-axis). (B) The kinetics of ligand uptake for a modeled ligand is simulated for different recycling and lysosomal degradation rates given in panel (A)-dotted line, representing the rates associated with LRBA-deficient cells. The results are obtained numerically from model (7) for an *in silico* cell culture experiment with a cell count of 10^6^ cells/ml, and with initial extracellular ligand concentration of L_p,0_ = 100 nM.

## Discussion

In the cell-extrinsic regulatory function of CTLA4, an efficient depletion of costimulatory ligands (CD80/CD86) via transendocytosis relies on the trafficking features of CTLA4. In this study, an intracellular trafficking model is constructed by taking into account the essential sub-processes involved in the constitutive trafficking of transmembrane proteins. Additional complexities are added to the model in order to capture the experimental protocols for characterizing the kinetics of CTLA4 molecules. We derived analytic solutions of the model for the different experimental protocols, which enabled us to estimate all CTLA4 trafficking parameters (see **Table 1**). The analytic solutions derived from the mathematical models revealed the explicit dependencies of experimental measurements on each of the trafficking parameters. For example, these explicit solutions verified that the distribution (equation (2)) and the observed kinetics of CTLA4 (equations (20), (24), (28), (36) and (37)) are independent of the absolute value for the rate of protein synthesis. Therefore, cells with different steady state levels of CTLA4 synthesis would still exhibit similar CTLA4 kinetics.

CTLA4 molecules show an unusual distribution in the cell, with a higher fraction of total CTLA4 being stored in the cytoplasm in the flow equilibrium. The estimated parameters confirmed the expectation that this distribution results from very fast trafficking rates. The ratio of internalization to recycling rates is in the range of ≈14.

A large storage of CTLA4 in the cytoplasm in combination with fast recycling provides a temporary compensation for surface CTLA4 loss when protein synthesis is interrupted (see surface unlabeled CTLA4 in **Figure 3C** and **Figure 4**). The rate of CTLA4 molecules newly expressed on the cell surface depends on both the CTLA4 synthesis and the amount of the recycled free CTLA4 (see equation (1) and (6)). Together with the speed of internalization (ln2/*k_i_* = 2.4 minutes), these processes determine how rapid antibodies/ligands are removed from the extracellular medium or the opposing cell.

According to the model, the flow of free CTLA4 molecules to the plasma membrane is highest at the time of exposure to antibodies and decreases within 2.5 hours (see **Figure 8B**). During this period of time, the pool of cytoplasmic free CTLA4 molecules is exhausted and replaced by an antibody labeled pool (see **Figure 2**). Consequently, the rate of antibody removal from the medium is decreased and limited by the binding to newly synthesized CTLA4 molecules. Viewed in the context of the transendocytosis model, this result suggests that efficient ligand capture may be achieved when CTLA4 expressing T cells remain bound to a single APC for a short duration (less than 2.5 hours). The T cell experiences a refractory time during which the depleted cytoplasmic free CTLA4 is restored and the internalized ligands are degraded. This limits the efficiency of CTLA4 expressing T cells in removing target molecules from multiple subsequent APCs. In a two-photon imaging study, it was shown that regulatory T cells (Treg), which constitutively express CTLA4 [33, 34], make transient interactions with lymph node resident dendritic cells (DC) with an average duration of 3.3 minutes, and up to 40 minutes with migratory activated DCs [35].

According to model (1), these contact durations correspond to a reduction of 8%–65% of the cytoplasmic free CTLA4 (see equation (17a)). It takes 11.5 hours for a CTLA4 depleted cell to increase the cytoplasmic free CTLA4 to 95% of its maximum value (data not shown). This duration decreases to approximately 10, 7 and 4.5 hours if the cytoplasmic CTLA4 pool is depleted by 65%, 25% and 8%, respectively. Therefore, upon a shorter exposure of CTLA4 expressing cells to ligands, a shorter refractory time is expected. This suggests that a full depletion is not achieved within a single interaction but that even short interactions of a few minutes would require a refractory time of a few hours before a CTLA4-expressing Treg would be expected to regain full power.

The functional limitation induced by the depletion of the cytoplasmic free CTLA4 and the required refractory time can be overcome if CTLA4 molecules return to the recycling pool following ligand binding (CTLA4 recovery). In this scenario, CTLA4 molecules bound to their target would dissociate from the target right after internalization and reenter the pool of free cytosolic CTLA4 available for recycling (see **Figure 12**). The CTLA4-target off-rate would control the rate of CTLA4 recovery. On the one hand, a low off-rate would limit CTLA4 recovery. On the other hand, a low off-rate increases the life time of each CTLA4:ligand complex on the cell surface and, by this, the probability of transendocytosis. Therefore, a subtle balance between ligand binding reaction and CTLA4 trafficking is needed for maintenance of efficient transendocytosis in a stable CTLA4^+^ T-APC interaction. The model predicts how the kinetics of ligand uptake is altered by binding affinities (**Figure 10**), and trafficking parameters (**Figure 11**).

CTLA4 occupancy on the cell surface depends on both, concentration and affinity to the ligand (see **Figure 10C**). However, in the cytoplasm, the dependency on ligand concentration saturates above a critical concentration (**Figure 10D**). This has implications for therapies that block CTLA4 with therapeutic ligands (CTLA4 antibodies): It is not possible to increase the efficiency of CTLA4 blockade by higher concentration of the ligand. In contrast, modulations of the ligand off-rate may change the efficiency of the drug. Additionally, it has to be taken into account that ligand uptake is optimally fast for a range of off-rate values, and decreasing off-rate values may coincide with a facilitated ligand removal and shorter drug effect (compare **Figures 10B** and **D**).

For modeled ligands with off-rates in the range of CD80/CD86 off-rates, cytoplasmic free CTLA4 is not reduced, irrespective of the ligand concentration (see vertical dotted-lines in **Figures 10D**). This suggests that the flow of free CTLA4 to the surface is not affected by complex formation with the natural ligand and complex internalization, and, consequently, ligand uptake keeps working at maximum capacity. However, the maximum capacity of keeping endocytosed ligands and intracellular degradation of ligands may be limited, which are not considered in our model.

In order to observe CTLA4 molecules in experiments, anti-CTLA4 antibodies were used. These molecules distribute in the medium, which allows them to bind to CTLA4 at any location of the cell surface. This three-dimensional access to CTLA4 by free antibodies in the medium is different from membrane-bound ligands expressed by APCs. In the context of T-APC interactions, upon cytoskeletal polarization of T cells towards APCs, cytoplasmic CTLA4 is translocated to the immunological synapse, which may facilitate CTLA4 exocytosis and increase CTLA4 surface density in the two-dimensional interaction face [17, 36]. This could result in a significantly higher binding rate of ligands and CTLA4 than in solution. In this regard, antibodies should have much higher affinities than the natural ligands in order to compensate for the enhanced binding in a synapse. Furthermore, continued endocytosis of CTLA4 into clathrin-coated pits at the immunological synapse could result in a faster rate of internalization. However, since T-APC interactions are discrete events, which depend on the chemokine environment, cell density and motility, transendocytosis does not occur continuously. In contrast, anti-CTLA4 antibodies in the medium can be constantly removed and degraded via CTLA4 (**Figure 12**), which may balance the more efficient binding within synapses to some degree.

The presented and experimentally validated modeling framework provides new insights into the control of immune responses by transendocytosis. Having established such a model, important biological questions related to immune regulation and autoimmune disease can be addressed. For example there are two natural ligands CD80 and CD86 with different binding affinities for CTLA4. The impact of different binding parameters and hypothetical changes in CTLA4 behavior resulting from this can be further explored. Moreover, it is already evident that mutations in CTLA4 [37] and most likely in its ligands will occur in the context of diseases. Such mutations may affect the binding affinity with unknown consequences. Accordingly, as we refine our model with experimental data it would become possible to predict the impact of such changes on immune function.

**Figure 12:**
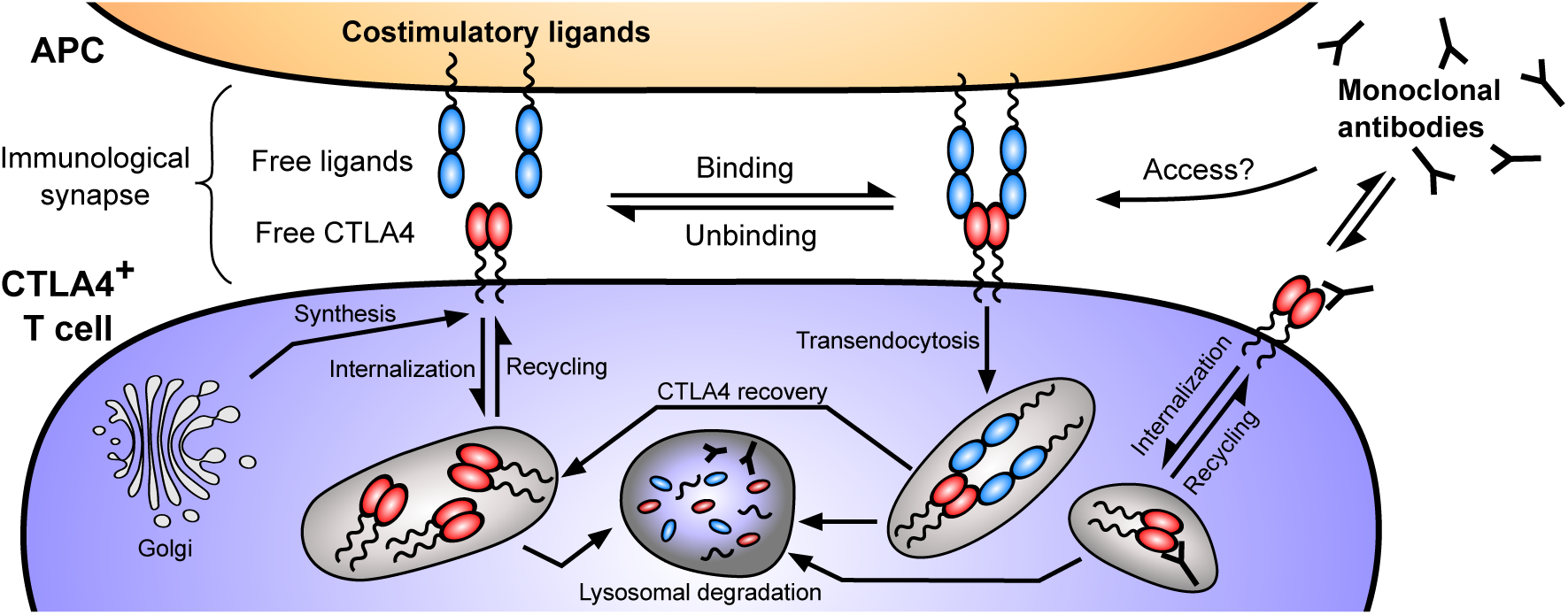
Free versus membrane-bound binding to CTLA4. Binding of anti-CTLA4 antibodies in the medium can continually occur because of three-dimensional access to surface free CTLA4 molecules. In contrast, binding of free CTLA4 and the membrane-bound ligands are restricted to the interaction of T cells and APCs. Despite this restriction, upon formation of the immunological synapse and cytoskeletal polarization of T cells toward APCs, a more efficient binding reaction in the two-dimensional T-APC interaction surface would occur. Due to the directed exocytosis of newly synthesized CTLA4 molecules towards the immunological synapse, and very tight and unique structure of the interaction surface, free anti-CTLA4 molecules may not access to this area. However, the effect of such a restriction, if true, would highly depend on the duration of T-APC interaction. A higher off-rate of natural ligands compared to anti-CTLA4 antibodies may allow the recovery of CTLA4 molecules, which bring their target molecules in the cytoplasm. In such event, recycling of costimulatory ligands in the recipient cell would not occur.

## Materials and methods

### CTLA4 internalisation assay

Chinese Hamster (*Cricetulus griseus*) Ovary (CHO) cells, expressing the virally transduced (MP-71 retroviral vector containing) human CTLA4 (UniProtKB accession: CTLA4 HUMAN) were placed in FACS tubes [SARSTEDT] at 1.5×10^6^ cell/mL density (150 μL) before being placed on ice. The cells were washed in 200 μL PBS‑ buffer (PBS pH 7.4 [Life Technologies] and 2% w/v Albumine Bovine Serum (BSA) [ACROS Organics]) by spinning cells at 500 g for 5 minutes. The surface CTLA4 was stained using Phycoerythrin (PE) fluorophore-conjugated mouse anti-human CTLA4 antibody (PE-CTLA4) [BD Pharmingen] at 1:100 dilution for 20 minutes on ice. After removing the excess antibody and washing the cells with ice cold PBS‑ buffer, the cells were placed at 37°C water bath for various duration, allowing CTLA4 internalisation from the CHO cell surface, before being placed on ice and fixed with 3% (v/v) paraformaldehyde (PFA) [Agar Scientific] for 15 minutes. The cells were washed and the surface CTLA4 was stained with the Alexa Fluor 647 conjugated chicken anti-mouse secondary antibody [ThermoFisher Scientific] at 1:1000 dilution for 30 minutes on ice before being washed and run through BD LSRFortessa (5L SORP) flow cytometer using yellow-green 561nm (586/15 band pass filter) and red 640nm (670/14 band pass filter) lasers for PE and Alexa Fluor 647 fluorophores, respectively. The data were recorded using BD FACSDiva Software (Version 6.2) and the emission spectra compensated mean fluorescence intensity (MFI) of each fluorophore was obtained using FlowJo 10.0.8r1 software.

### CTLA4 molecular counting

Quantum Simply Cellular anti-Mouse IgG kit [Bangs Laboratories, Inc.] were used according to the manufacturer’s instructions in order to quantify the number of virally transduced human CTLA4 expressed in CHO cells. Briefly, beads 1-4, which are of known size and antibody binding capacity, were stained with varying amounts of PE-CTLA4 for 30 minutes on ice. The beads were washed in media (DMEM [Life Technologies], 10% FBS [Labtech], 2 mM L-Glutamine [Life Technologies], 1x Penicillin-Streptomycin [Life Technologies]) by spinning at 1060 g for 5 minutes before their median fluorescence intensity (MFI) were obtained using BD LSRFortessa flow cytometer using the yellow-green 561nm (586/15 band pass filter) laser at a set voltage. Bead saturation levels with PE-CTLA4 antibody were assessed using the FlowJo 10.0.8r1 software and the best MFI for each bead was plotted in order to establish a calibration curve. In parallel to the above, various number of CHO cells expressing human CTLA4 were fixed with 3% (v/v) PFA [Agar Scientific] for 15 minutes on ice before being washed in media and permeabilised with 0.1% (v/v) saponin in media. The cells were then stained in an identical manner to the beads and ran through BD LSRFortessa flow cytometer using identical settings and voltages as the beads. The obtained CHO cell MFIs were then compared against the calibration curve in order to obtain the number of CTLA4 molecules per CHO cell.

### CTLA4 staining model

A typical staining protocol of a protein with natural trafficking could be represented by a nonlinear model schematically illustrated in **Figure 13**. In this model, it is assumed that free CTLA4 molecules are synthesized by a constant rate *σ_i_* and directly exocytosed to the plasma membrane (*R_p_(t)*). The free surface CTLA4 molecules can bind to labeling antibodies (*L_p_(t)*) and form CTLA4:antibody complex 
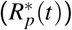
 which can be detected by flow cytometry or confocal microscopy. The extracellular staining reaction 
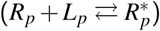
 occurs with a constant binding rate *k_L_* and unbinding rate 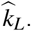 Free and labeled CTLA4 molecules on the plasma membrane (*R_p_*(*t*) and 
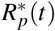
, respectively) are internalized with constant rates *k_i_* and *k_i_*^*^, respectively. Once the labeled CTLA4 molecules are in the cytoplasm (*R_c_*^*^(*t*)), free labeling antibodies (*L_c_*(*t*)) appear in the cytoplasm via unbinding of the complex to the reactants, which can further bind to free cytoplasmic CTLA4 molecules *R_c_*(*t*) 
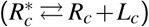
. The cytoplasmic free and labeled CTLA4 molecules are recycled to the plasma membrane with constant rates *k_r_* and 
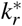
, respectively. Note that in this model, internalization is assumed to be the only pathway by which labeling antibodies could enter the cell cytoplasm from the extracellular environment. Given these assumptions, the CTLA4 staining model can be described by the following set of ODEs

**Figure 13:**
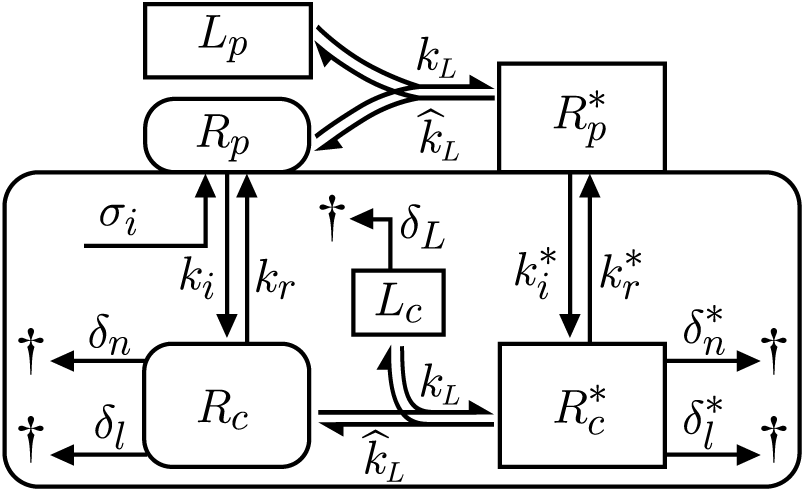
CTLA4 staining model.

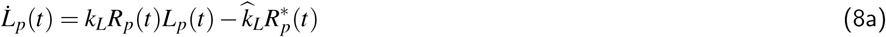

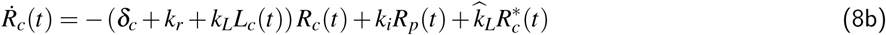

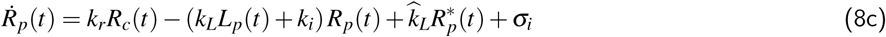

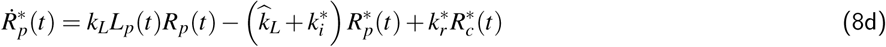

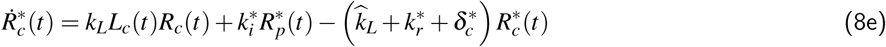

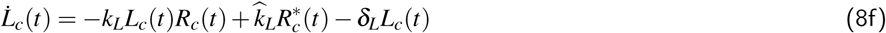

The nonlinearity of this model arises from the forward reaction of extracellular and intracellular CTLA4 labeling.

### Reduction of the CTLA4 staining model

We use the following assumptions to reduce the staining model (8): (A) *L_p_* is constant and sufficiently large, (B) *k_L_* is sufficiently larger than other reaction rates, 
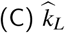
 is smaller than the other rates and is approximated to zero for the duration of the experiment, and (D) 
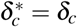
, 
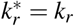
 and 
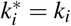
. Assumptions (A) and (B) imply that the labeling of the surface CTLA4 molecules (*R_p_*(*t*)) is at quasi steady state. By setting *Ṙ_p_*(*t*)^˙^ = 0 in equation (8c), we have

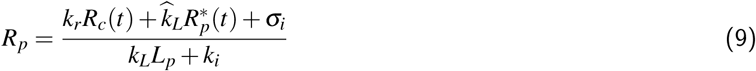

Since the cytoplasmic staining antibody *L_c_*(*t*) appears during the unbinding process of internalized labeled CTLA4 
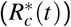
, its value at the starting time of the labeling process (*L*_*c*,0_) is zero. Then, by employing assumption (C), we evaluate equation (8f) with 
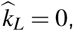

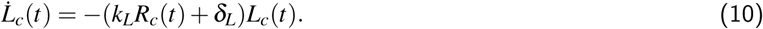

Therefore, *L_c_*(*t*) remains at its equilibrium point, i.e.

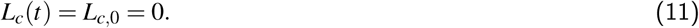

Next, we substitute (9) and (11) in (8b),

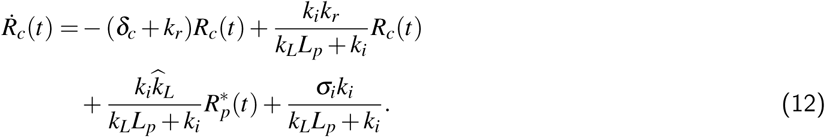

According to assumptions (A) and (B), we extract *k_L_L_p_*≫*k_i_k_r_*,

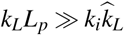
 and *k_L_L_p_*≫σ_*i*_*k_i_*. Therefore, the ratios in equation (12) can be approximated to zero, and equation (12) can be simplified to

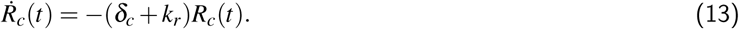

Next, we evaluate equation (8d) according to equations (9) and (11) and assumptions (C) and (D),

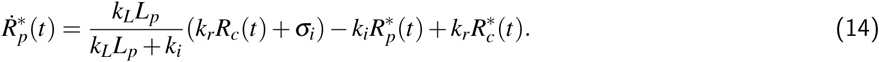

From assumptions (A) and (B), we extract *k_L_L_p_*≫*k_i_*. Therefore, 
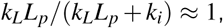
 and equation (14) can be simplified to

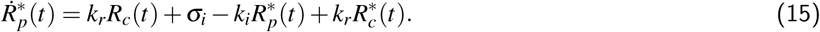

Finally, we evaluate (8e) according to (9), (11) and assumptions (C) and (D), and obtain

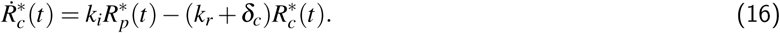

The reduced model is given in equation (3), and is shown schematically in **Figure 1**.

With initial conditions 
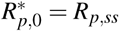
, *R*_*p*,0_^*^ = *R_p,ss_* (see equation (2)) and 
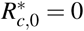
, the solution of the model (3) is

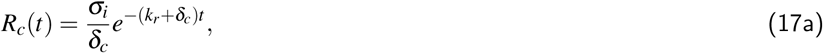

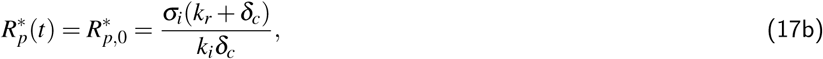

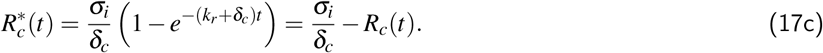

#### CTLA4 internalization model

The theoretical amount of initially labeled CTLA4 molecules on the cell surface 
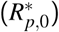
 is the same as equation (2), i.e. 
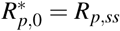
. After washing, the kinetics of labeled CTLA4 molecules on the plasma membrane follows equation (4). From the ODEs in (3c) and (4), the number of labeled CTLA4 molecules can be obtained as

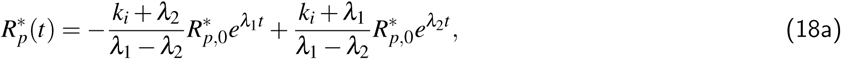

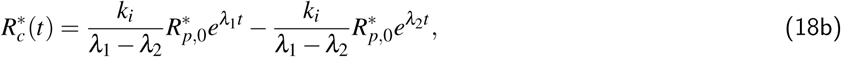

Where λ_1,2_ belongs to the set of eigenvalues of the CTLA4 staining model (3),

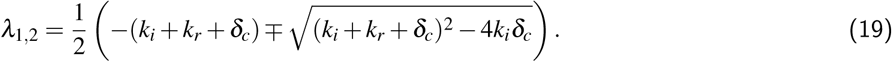

For biologically relevant parameters (with positive values), λ_1,2_ are negative.

For the purpose of parameters estimation, the fraction *Y*_int_ of initially labeled CTLA4 molecules that remained on or recycled to the cell surface is of interest (see **Figure 3A**), which is equivalent to

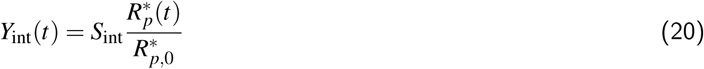

Where *S*_int_ is an unknown scaling factor that allows estimation of the initial data point, and is fitted during parameters estimation.

#### Block of CTLA4 synthesis model

Consider the model (1) with imposing σ_*i*_ = 0 during the presence of CHX. At the moment of adding CHX (*t* = 0), the amount of CTLA4 molecules (*R*_*p*,0_ and *R*_*c*,0_) is equal to the steady states of the model (1), given in equation (2). With these initial conditions, the CTLA4 levels in the presence of CHX vary according to the following functions where are the constant coefficients and λ_1,2_ are given in equation (19).

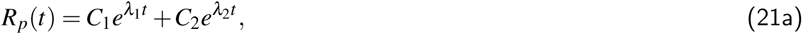

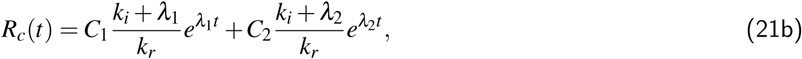

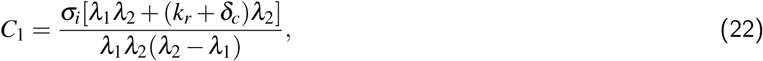

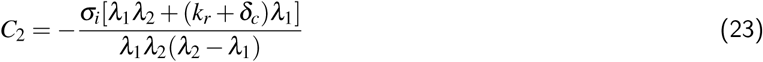

The theoretical value that corresponds to the experimental measurement is

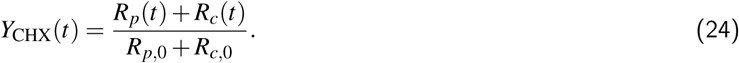

Note that in equation (24), the rate of protein synthesis (σ_*i*_) does not appear.

#### Block of lysosomal degradation

From the solution of the CTLA4 staining model (3) given in equation (17), we obtain the amount of labeled CTLA4 molecules after 60 minutes on the plasma membrane 
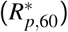
 and in the cytoplasm 
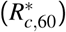
. Note that in this stage of p 60c 60 the experiment, we have δ_*c*_ = δ_*l*_ + δ_*n*_. After washing cells, labeled CTLA4 molecules change over time according to equations (3c) and (4) with the following solutions where are the constant coefficients and the values for λ_1,2_ have the same form as in equation (19), but with different rates of cytoplasmic degradation (δ_*c*_) reflecting the different experimental condition; in the presence of NH_4_Cl in the medium, we have δ_c_ = δ_*n*_ whereas in the control experiment δ_*c*_ = δ_*l*_ + δ_*n*_ holds.

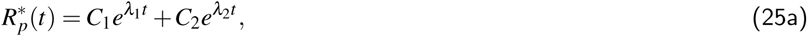

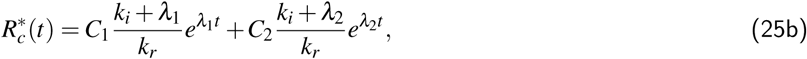

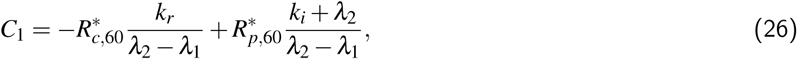

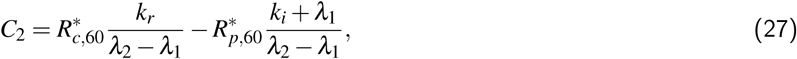

Experimentally measured MFI values for the lysosomal-block experiment (*Y_NH_*_4*Cl*_(*t*)) and the control experiment (*Y*_CTR_(*t*)) are correlated with the subcellular labeled CTLA4 molecules Note that in these observers, the rate of lysosomal degradation is different.

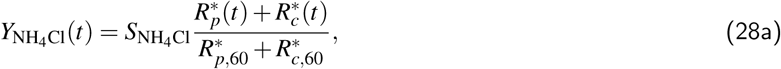

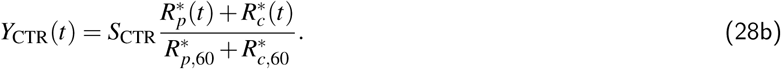

### CTLA4 recycling model

Following the recycling experiment, the corresponding model is constructed in two steps. This model is illustrated schematically in **Figure 14A-B**. During the first step of staining with green primary antibodies, a subset of CTLA4 molecules was labeled at the cell surface and subsequently internalized. The labeled CTLA4 on the plasma membrane and in cytoplasm are represented by 
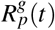
 and 
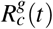
, respectively. From the staining model (3) and steady state solutions in equations (17b) and (17c), we obtain the amount of green labeled CTLA4 molecules 
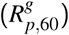
.

Then in experiment, green-labeled CTLA4 molecules on the cell surface 
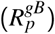
 are blocked on ice, which imposes an initial value of 
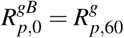
 equal to the amount of surface green labeled CTLA4 molecules after the first staining step. The cytoplasmic blocked pool 
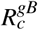
 changes from its initial value 
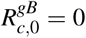
 only after raising the temprature to 37°C.

**Figure 14.**
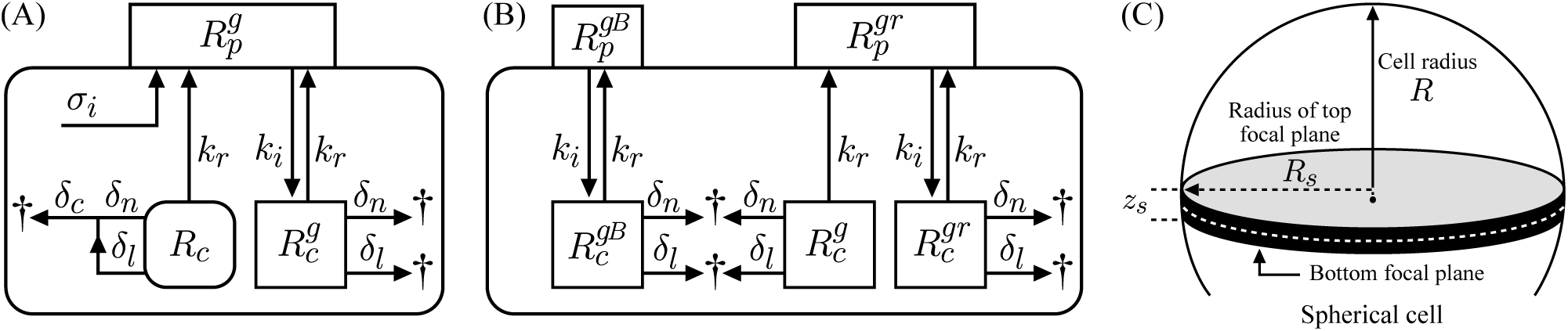
4: CTLA4 recycling model. The recycling model is developed for two staining steps. (A) Schematic of the first step staining with Alexa488-conjugated anti-CTLA4 (green). (B) Schematic of the second stage staining with Alexa555-conjugated anti-human IgG secondary antibody (red), during which the recycling green-labeled CTLA4 molecules occurs after initially blocking the green-labeled CTLA4 molecules on plasma membrane (on ice). Note that the amount of green-labeled CTLA4 molecules after 1 hour in the first stage is used as the initial conditions for the second stage. (C) Visible area and volume of a spherical cell in confocal microscopy.

Next, after washing and raising the temperature, the amounts of red-green labeled CTLA4 molecules on the plasma membrane 
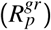
 and in the cytoplasm 
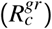
 change from their initial values 
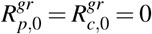
. During this second staining step, the amount of different CTLA4 pools relies on the following dynamical model

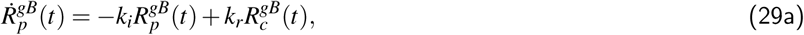

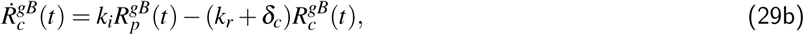

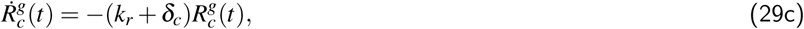

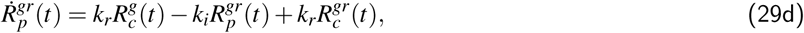

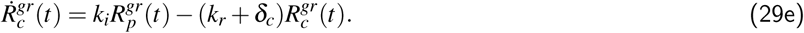

The dynamics of unlabeled CTLA4 molecules is the same as in model (1). The submodel consisting of equations (29a) and (29b) is equivalent to the internalization experiment, and the submodel consisting of equations (29c-e) is equivalent to the homogeneous version of the staining model (3). The solution of model (29) with the given initial conditions is the following where λ_1,2,3_ are are the eigenvalues of staining model (3). The values for λ_1,2_ are given in equation (19) and λ_3_ = (*k_r_* + δ_*c*_).

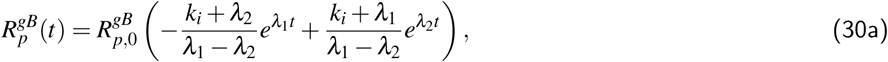

x
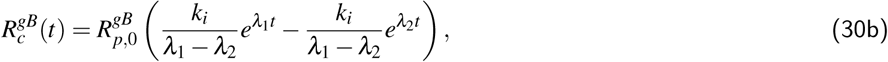

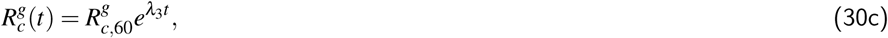

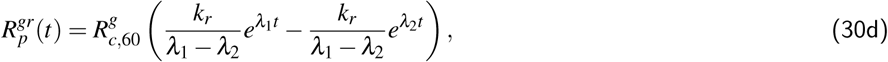

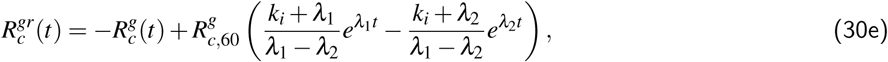

All the green‑ and red-labeled CTLA4 molecules were quantified by confocal microscopy. Note that by confocal microscopy, only a focus plane is visible, and only a fraction of CTLA4 molecules is observable. Therefore, this fraction has to be considered in the theoretical values. In general, by considering a visible area *A_vis_* and a visible volume *V_vis_* of a spherical cell with area *A* and volume *V*, the following fractions of labeled CTLA4 molecules on the plasma membrane 
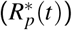
 and in the cytoplasm 
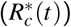
 can be observed:

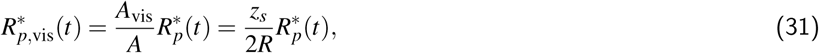

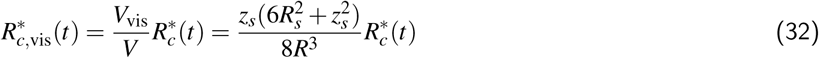

Where *R* is the radius of the spherical cell, *z_s_* is the thickness of the cell slice that is visible by the confocal microscopy and

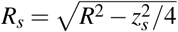
 is the radius of top focal plane (see **Figure 14C**). Theoretical numbers of green‑ and red-labeled CTLA4 molecules that are visible by confocal microscopy, denoted by *G*(*t*) and *R*(*t*) respectively, are

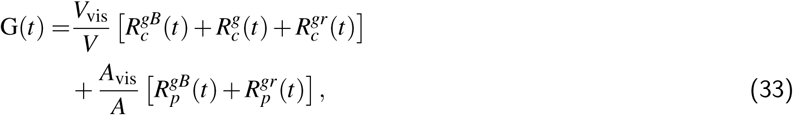

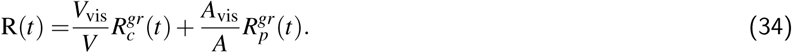

These theoretical values are assumed to be proportional to the measured MFI values. For parameter estimation, the same type of normalization as applied to the experimental data has to be considered for the theoretical values. These normalizations are

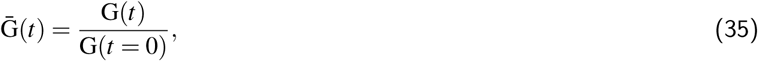

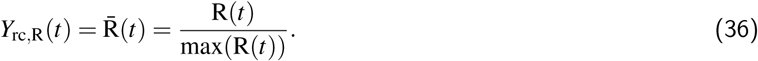

Note that in (35), the theoretical and experimental values for *t* = 0 are equal to 1. Therefore, normalization imposes a zero estimation error for the initial data point of the green-labeled pool. To avoid this, we impose a scaling factor *S*_rc_ which allows an estimation of the initial data point. Furthermore, nonzero value of red fluorescent intensity in **Figure 6A** at *t* = 0 indicates a delay between the starting time of the recycling process and the measurement. Therefore, we define a shifting time (Δ*t*_rc_) in the model. With these considerations, the theoretical value for the green fluorescent measurement to be used for parameter estimation reads

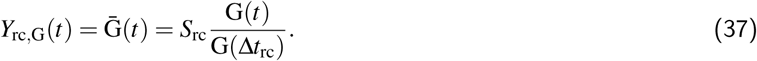

Since we considered the time shift in the experimental values and the red fluorescent measurements are normalized to the maximum value but not to the initial data point, the theoretical value (35) does not need any modification and is suitable for parameter estimation. Note that in equations (36) and (37), the rate of protein synthesis (σ_i_) does not appear. The fitting result is shown in **Figure 6A
**.

### Parameter estimation

The ligand-independent CTLA4 trafficking model in its simplest form in equation (1) has 5 unknown parameters. There are 5 additional parameters that are concerned with the types of experimental protocols, and aimed to handle uncertainties. The list of parameters and associated dimensions are given in **Table 1**. In order to find an optimal parameters set that fits the modeling results to the experimental observations, the following cost (objective) function *E*(θ) is minimized

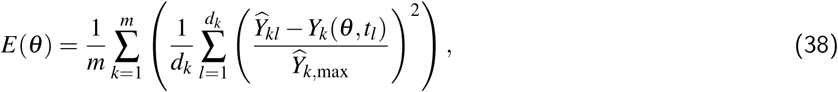

Where *Ŷ_kl_* is the observed value from *k*-th experiment at time *t* = *t*_*l*_ and *Y*_*k*_(θ *t*_*l*_) is corresponding theoretical value (observer) obtained from the model with parameter set θ. *d*_*k*_ is the number of data points in *k*-th experiment, *Ŷ*_*k*,max_ is the maximum observed experimental value in *k*-th experiment, and *m* is the number of independent experiments. We used a differential evolution algorithm to minimize the cost function in equation (38). The estimated parameter values are given in **Table 1**.

## Acknowledgments

S. K. acknowledges the support of the German Federal Ministry of Education and Research (BMBF) for the eMED project SYSIMIT. B. R. was supported by UK Biotechnology and Biological Sciences Research Council (BBSRC). P. A. R. and M. M.-H. were supported by the Human Frontier Science Program (RGP0033/2015). M. M.-H was supported by iMed – the Helmholtz Initiative on Personalized Medicine. The authors wish to thank Sebastian Binder and Ghazal Montaseri for revising the manuscript.

